# Multidimensional analysis of cortical interneuron synaptic features reveals underlying synaptic heterogeneity

**DOI:** 10.1101/2024.03.22.586340

**Authors:** Patrick D. Dummer, Dylan I. Lee, Sakib Hossain, Runsheng Wang, Andre Evard, Gabriel Newman, Claire Ho, Casey M. Schneider-Mizell, Vilas Menon, Edmund Au

## Abstract

Cortical interneurons represent a diverse set of neuronal subtypes characterized in part by their striking degree of synaptic specificity. However, little is known about the extent of synaptic diversity because of the lack of unbiased methods to extract synaptic features among interneuron subtypes. Here, we develop an approach to aggregate image features from fluorescent confocal images of interneuron synapses and their post-synaptic targets, in order to characterize the heterogeneity of synapses at fine scale. We started by training a model that recognizes pre- and post-synaptic compartments and then determines the target of each genetically-identified interneuron synapse in vitro and in vivo. Our model extracts hundreds of spatial and intensity features from each analyzed synapse, constructing a multidimensional data set, consisting of millions of synapses, which allowed us to perform an unsupervised analysis on this dataset, uncovering novel synaptic subgroups. The subgroups were spatially distributed in a highly structured manner that revealed the local underlying topology of the postsynaptic environment. Dendrite-targeting subgroups were clustered onto subdomains of the dendrite along the proximal to distal axis. Soma-targeting subgroups were enriched onto different postsynaptic cell types. We also find that the two main subclasses of interneurons, basket cells and somatostatin interneurons, utilize distinct strategies to enact inhibitory coverage. Thus, our analysis of multidimensional synaptic features establishes a conceptual framework for studying interneuron synaptic diversity.

## Introduction

Cortical interneurons are a highly diverse cell population and our understanding of that diversity has accelerated as a result of high throughput transcriptomic-based studies in recent years ^1–3^. A knowledge gap exists, however, regarding the diversity of cortical interneuron synapses. While we know that interneurons target different compartments of the postsynaptic neuron – dendrite, soma and axon initial segment – it is not clear whether further heterogeneity exists. A growing body of evidence links cortical interneuron dysfunction to neuropsychiatric illness, a constellation of disorders typified by defects in synaptic function ^4–8^. Furthermore, an enhanced understanding of interneuron synaptic diversity is key to linking transcriptomic cellular identity and the diverse functions of interneurons in the cortex. Thus, a framework for categorizing interneuron synaptic heterogeneity would be a powerful tool for revealing interneuron function in the healthy brain and how interneuron dysfunction contributes to disease etiology.

Our understanding of interneuron synaptic diversity is hampered by a lack of high throughput methodologies that can extract multidimensional data from individual synapses the way that single cell RNA-seq (scRNA-seq) data allows for the analysis cellular diversity. To address this, we reasoned that synapses are, at their core, a spatial apposition between 2 neurons. Thus, an image-based spatial analysis between pre- and post-synaptic compartments might allow us to extract biologically meaningful features from synapses. These features, in turn, could be analyzed at a population level to reveal underlying interneuron synaptic heterogeneity.

To develop our imaged-based approach, we made use of 2 well-studied principles in interneuron synaptic connectivity. First, interneurons form connections onto different cellular compartments of their postsynaptic partners – dendrite, soma and axon initial segment (AIS). Second, different interneuron subclasses exhibit strong target preferences for each compartment. Somatostatin-positive (Sst) interneurons prefer to target dendrite ^9,10^, basket cells (BCs) largely target soma and proximal dendrite ^11,12^, and chandelier cells (ChCs) have a strong preference for AIS ^13,14^. Interneuron subclasses form repetitive circuit motifs throughout the cortex ^15^. This repetitive connectivity lends itself to studying interneuron synapses at a population level where any given area of the cortex can provide a representative sample of all types of interneuron synapses.

Given these prior findings on interneuron synapse diversity, we hypothesized that representative imaged fields of interneuron synapses could serve as starting material for a computer model that was trained to perform a relatively simple task: assigning the target compartment for every synapse imaged. In the process of performing this supervised classification task, the model would extract multiple features from each synapse analyzed, based solely on local image statistics, and not sub-selected or refined specifically for the classification task. Thus, the model would concurrently generate a rich, multidimensional dataset that could reveal underlying synaptic heterogeneity that is invisible to the human eye.

In this study, we start with standard 4-channel confocal images of interneuron synapses and their postsynaptic compartments. From this, we devised an automated, high-throughput method to extract spatial and intensity metrics between pre- and post-synaptic features that were used to assign targets for each synapse at the scale of hundreds of thousands of synapses per analysis. We then performed an unsupervised cluster analysis based on synaptic features, which identified multiple subgroups of dendrite- and soma-targeting synapses. Spatial analysis of synaptic subgroups revealed novel insights into interneuron synaptic biology.

## Results

### Labeling Pre- and Post-synaptic Compartments

Interneurons target different compartments of the postsynaptic neuron – dendrite, soma and AIS. To establish an image-based system that can automatically assign a target for each synapse, we first validated an immunohistochemical strategy to label target compartments in both cortical sections as well as in dissociated cortical culture (Figure 1A). To label soma and proximal dendrite, we used a combination of antibodies against voltage-gated potassium channels, Kv_2.1_ and Kv_2.2_ (collectively Kv2), combined in the same channel ^16^. To label axon initial segment, we used an antibody against Ankyrin G ^17^. Labeling dendrite was more challenging, and our attempts with MAP2 were unsuccessful because of uneven signal intensity and the fact that MAP2 only labels dendritic shaft (Supplemental Figure 1). Although inhibitory input is primarily onto dendritic shaft, a sizeable portion innervates spine and neck ^18,19^. As an alternative, we opted for a negative selection approach: we used an antibody against the pan-inhibitory postsynaptic scaffold protein, gephyrin and then relied on the absence of Kv2 and Ankyrin G to define the dendritic target compartment.

**Figure 1.**
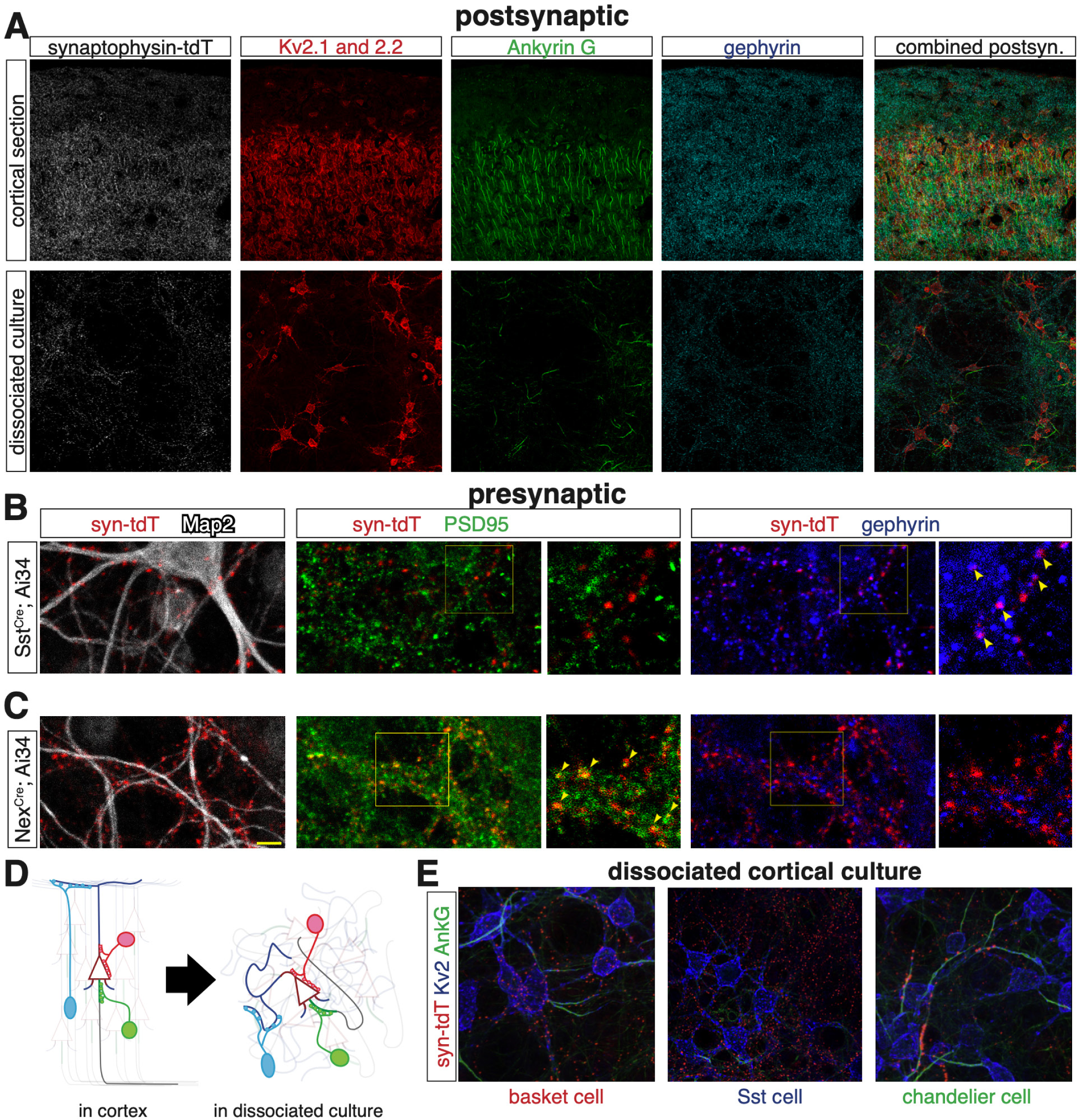
Labeling target compartments and presynaptic puncta for image-based analysis system. (A) Immmunohistochemical labeling of postsynaptic target compartments in (top row) cortical sections and (bottom row) dissociated cortical culture. Target compartments labeled: soma and proximal dendrite (Kv_2.1_ and Kv_2.2_ combined, Kv2), axon initial segment (Ankyrin G), dendrite (gephyrin). (B, C) Validation of genetic labeling of presynaptic synaptophysin-tdTomato+ puncta. (B) Pan-MGE lineage interneuron Cre driver, Nkx2-1^Cre^; Ai34 presynaptic puncta counter labeled with inhibitory synaptic marker (gephyrin, blue) and excitatory synaptic marker (PSD95, green). (C) Pan-pyramidal cell Cre driver, Nex^Cre^; Ai34 presynaptic puncta counterlabeled with gephyrin (blue) and PSD95 (green). Yellow arrowheads indicate overlap/close apposition between syn-tdT puncta and markers. (D) Schematic showing canonical target preference for 3 interneuron subclasses: Sst interneuron, basket cell (BC) and chandelier cell (ChC). Right panel shows targeting in cortex, left panel shows target recapitulation in dissociated culture. (E) Dissociated cultures in which Sst, BC and ChCs were genetically-labeled with Ai34 reporter. Counterlabeled with Kv2 (blue) and Ankyrin G (green).

A second necessary component for our image-based system is to label presynaptic interneuron boutons. For this, we made use of a Cre-dependent conditional reporter allele (Ai34 ^20^) that expresses synaptophysin-tdTomato (syn-tdT) fusion protein. This allowed us to label presynaptic boutons with specificity achieved through various Cre driver lines. To validate the Ai34 reporter, we crossed it to Nkx2-1^Cre^, a pan MGE lineage interneuron Cre driver (Figure 1B), and Nex^Cre^, a pan pyramidal cell Cre driver (Figure 1C) and counterlabeled with inhibitory (Vgat and gephyrin) and excitatory (PSD95) synaptic markers. Consistent with our expectations, the vast majority of Nkx2-1^Cre^; Ai34 labeled syn-TdT puncta were co-labeled with Vgat (97%) and apposed to gephyrin puncta (96%) while not showing a close spatial relationship with PSD95. In contrast, Nex^Cre^; Ai34 labeled syn-tdT puncta were nearly all juxtaposed with PSD95 while not associating with gephyrin or Vgat.

Next, we crossed Ai34 to interneuron Cre driver lines and immunolabeled with postsynaptic markers for targets (Supplemental Figure 2A-C). As expected, syn-tdT puncta were enriched on Kv2-positive somata in Pv^Cre^ cortical sections, whereas in Sst^Cre^ cortical sections, syn-tdT was enriched with gephyrin puncta in the absence of Kv2. We did not observe syn-tdT puncta present in layer 1 in Pv^Cre^, but did so in Nkx^Cre^ and Sst^Cre^ cortical sections. This was as expected since Nkx^Cre^ and Sst^Cre^ label Sst interneurons that densely innervate pyramidal neuron distal dendrites.

Interneuron subclasses have strong subcellular target preferences – Sst interneurons for dendrite, BCs for soma and proximal dendrite and ChCs for AIS. We wished to label each subclass specifically to serve as a positive control for our approach. For Sst interneurons, we used Sst^Cre^. For BCs and ChCs, however, we needed an alternate approach since current Cre driver lines cannot distinguish between BCs and ChC (both of the parvalbumin-(Pv) positive lineage). It was previously shown that the Nkx2-1^CreERT2^ driver with an oral tamoxifen gavage at e17.5 labels ChCs with high enrichment in medial prefrontal cortex (pfc) ^21^. By late embryogenesis, interneuron production is primarily of the Pv lineage ^22^. We therefore hypothesized that while there was enrichment for ChCs in medial pfc, there might be areas of enrichment for BCs elsewhere in the cortex. To test this, we performed an e17.5 gavage on Nkx2-1^CreERT2^; Ai9 mice and analyzed at P40 (n=8 mice). We immunolabeled for Sst and Pv and found that vast majority of fate mapped interneurons were Pv-positive (94%) (Supplemental Figure 3A,B). We imaged multiple sections in the coronal plane throughout the rostral-caudal axis of cortex and developed an image analysis script to distinguish ChC cartridges from all other boutons (Supplemental Figure 3C). From this, we identified hotspots for labeled ChCs and areas where cartridges were absent, which we reasoned were areas enriched for BCs since nearly all labeled neurons were Pv-positive (Supplemental Figure 3D, E). To further confirm, we examined the apposition of syn-tdT puncta with postsynaptic targets in areas enriched for ChCs and BCs in Nkx2-1^CreERT2^; Ai34 mice, which confirmed our findings (Supplemental Figure 2D, E).

Previous work suggests that interneuron synaptic targeting relies upon a strong intrinsic component where pre- and post-synaptic molecules confer specificity in a lock and key fashion ^23–26^. Therefore, we tested whether interneurons in dissociated culture might retain synaptic specificity (Figure 1D). We generated dissociated cortical cultures from Sst^Cre^; Ai34 and Nkx2-1^CreERT2^; Ai34 pups in which we isolated regions enriched for BCs ChCs. Thus, 3 separate cultures were generated that labeled Sst, BCs or ChCs. We counterlabeled with postsynaptic target markers and found distinct differences in the target preferences for each of the 3 culture conditions that were consistent with subclass preferences for dendrite, soma and AIS, respectively (Figure 1E).

### Supervised Analysis of Interneuron Synaptic Targeting

Having established labeling strategies for postsynaptic targets and a Cre-dependent labeling strategy for subclass-specific presynaptic puncta, we proceeded with training a model to assign targets to all syn-tdT puncta. To do so, we decided to take an iterative approach: first train the computer on images obtained from dissociated culture where targeting cultures are effectively a single layer with lower synapse density, making training more definitive. This first round of training would then provide a sound foundation for a second round of training in cortical sections where the environment is more complex.

However, we noted that even in dissociated culture, the subcellular structures of interest were imperfectly represented by immunohistochemistry. This discrepancy proved difficult for a model to discern (Supplemental Figures 4 and 5) and necessitated that we train the model to define pre- and post-synaptic structures using features from multiple raw channels. To accomplish this, we utilized ilastik, a machine learning-based image analysis software program ^27,28^. Using ilastik we trained the model (pixel training) to define pre- and post-synaptic features through series of probability channels (Figure 2A-D) - 4 presynaptic and 7 postsynaptic channels, 11 channels in total. Combined with the original 4-channel confocal image, ilastik worked with 15 channels to define each image we capture.

**Figure 2.**
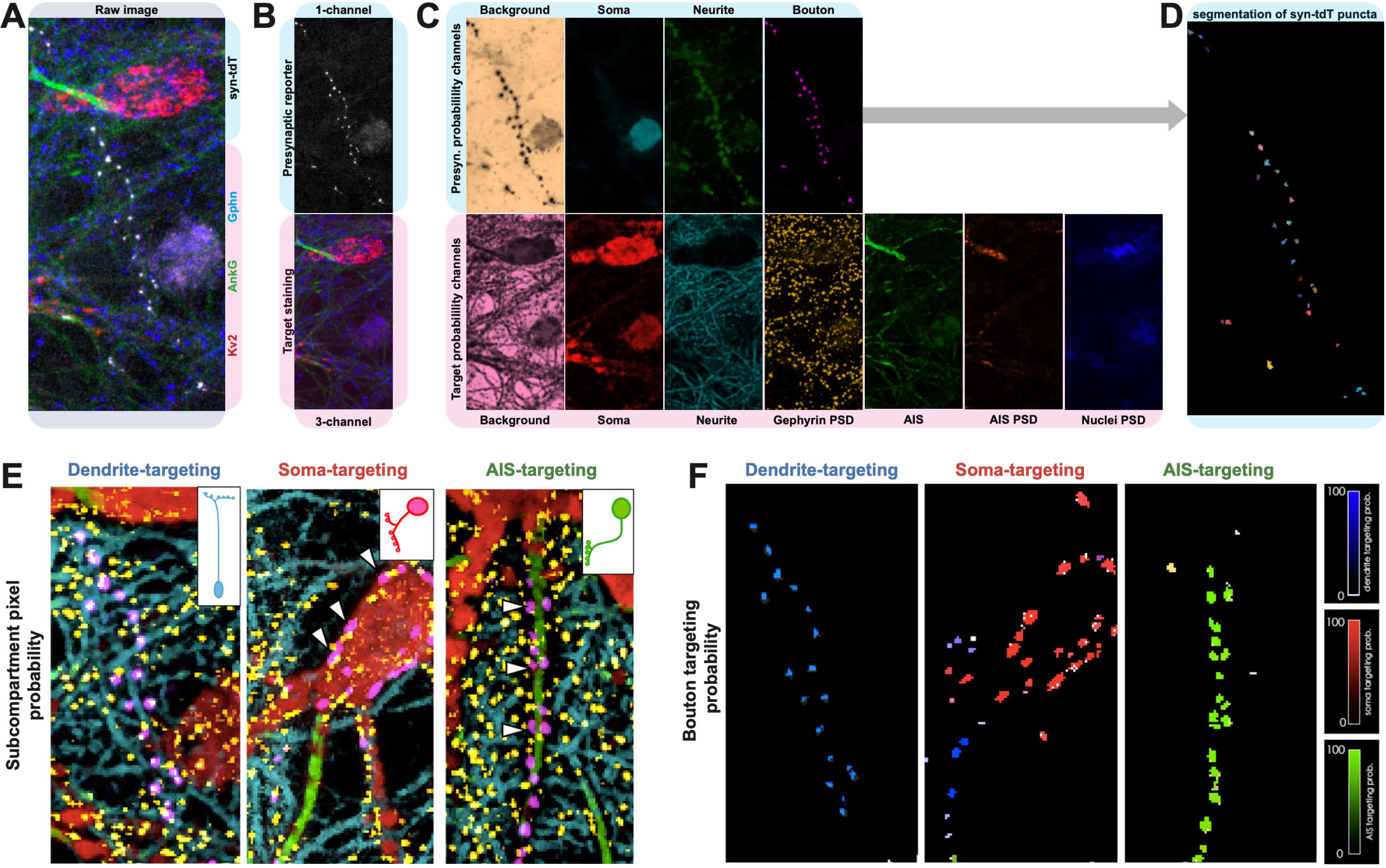
Overview of image-based model for assigning synaptic target to individual interneuron synaptic puncta. (A-D) Automated assignment of pre-(blue) and post-(pink) synaptic compartments to 11 probability channels. (A) Raw 4-channel confocal image for Ai34-label presynaptic puncta (white), Kv2 (red), Ankyrin G (green) and gephyrin (blue). (B) Raw channels split into pre-(top, blue) and post-(bottom, pink) synaptic compartments. (C) Raw pre- and post-synaptic channels split into probability channels. Presynaptic probability channels used to perform (D) object segmentation of syn-tdT presynaptic puncta. (E) Overlay of probability channels showing example images that were used for object training to define dendrite-, soma- and AIS-targeting synapses. (F) Target probability score assigned to each synapse, color coded according to heatmap (right) for each target compartment.

The 4 probability channels defining the presynaptic environment were used to perform object segmentation using FIJI (Figure 2E, Supplemental Figure 6A). This defines each syn-tdT puncta as separate objects to be individually assessed downstream. Each object was used for the next round of training (object training) in which we trained ilastik to recognize each syn-tdT puncta as targeting dendrite, soma, AIS, unknown or not a synaptic punctum (Figure 2F,G; Supplemental Figure 6B-D). This portion of the training was greatly aided by starting with image datasets obtained from dissociated monolayer culture where targeting is clearer, synapse density is lower (by approximately 10-fold) and the pre- and post-synaptic structures are effectively 2-dimensional.

After pixel and object training in dissociated culture, we performed a second round of pixel and object training on images of cortical sections. Since the model at that time was already reasonably proficient at recognizing targets, our second round of training in cortical sections mainly focused on slight correction and refinement for both pixel and object. In total, our pixel training dataset consisted of 297 images (213 culture, 84 cortical sections) with ∼180,000 training points. Our object training dataset consisted of 115 images (72 culture, 43 cortical section) with ∼9,000 training points.

Our model provided pixel probability maps and target assignment for syn-tdT puncta that appeared visually appropriate for both dissociated culture and cortical sections (Supplemental Figure 7) and the vast majority (> 98.5%) of puncta were definitively assigned a target. To test the model’s performance, we compare its results to manual human user scoring. We manually classified ∼5400 syn-tdT puncta that were randomly selected and not part of the training data set (4673 from culture, 796 from cortical sections). A user was trained on how to navigate ilastik and access the pixel probability maps generated from the image so that the user and the computer had access to the same information for assigning synapse targeting. Across all syn-tdT puncta, we found strong concordance (93.8% overall) between human and computer calls overall and that the accuracy was similar between boutons scored in culture (94.3%) versus cortical sections (90.3%) (Supplemental Figure 8).

### Interneuron Synaptic Targeting In Vivo and In Vitro

Having developed a model that reliably assigned cell compartmental targeting, we assessed synaptic targeting of interneuron subclasses in cortical sections and compared it with targeting in dissociated culture. Interneurons in dissociated culture lack multiple extrinsic cues for synaptic targeting: cortical structure, appropriate sensory input, network activity and paracrine factors produced in the cortical milieu. Thus, cultured interneurons must principally rely on intrinsic mechanisms to form appropriate synaptic connections and we could examine the extent to which cell-intrinsic mechanisms dictate interneuron target specificity by comparing synaptic targeting in culture versus in cortical sections.

First, we compared targeting in cortical sections between 4 interneuron Cre driver lines: 1) Nkx2-1^Cre^, which labels the entire MGE lineage (Sst, BC and ChC); 2) Pv^Cre^, which labels the Pv lineage (BC and ChC); Sst^Cre^, which labels the Sst lineage (Sst); and Nkkx2-1^CreERT2^, which labels primarily Pv-positive interneurons with BCs and ChCs enriched in different cortical regions (Figure 3A,B). We found significant differences in targeting between Nkx2-1 and Sst^Cre^ and between Sst^Cre^ and Pv^Cre^. Regions enriched for BCs and ChCs in Nkx2-1^CreERT2^ also showed strong and significant preferences for soma and AIS, respectively. While we observed an enrichment for AIS targeting in ChC enriched regions but it was not close to the near 100% AIS target fidelity thought to be exhibited by ChCs ^13,14^. We ascribe this discrepancy to the presence of non-ChC synapses in the same field and the relatively low number of synapses present in AIS-targeting cartridges versus other interneuron synapses, which served to dilute the AIS targeting numbers further. Additionally, we observed Sst and BCs forming targets onto targets other than dendrites and soma, which was expected and in line with proportion of ‘non-canonical’ synapses reported in prior studies ^9–11^.

**Figure 3.**
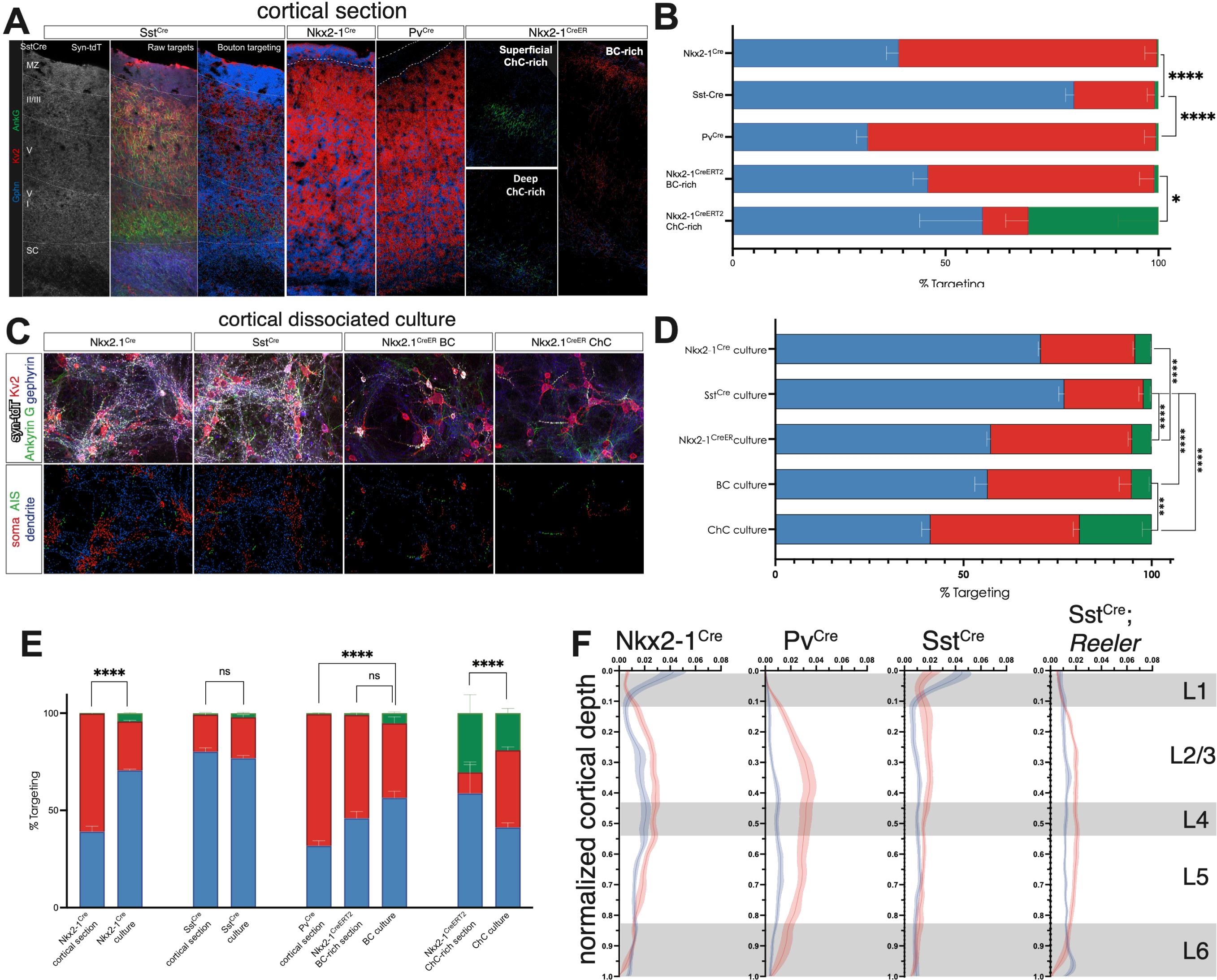
Quantification and comparison of interneuron targeting in cortical sections and in dissociated culture. (A) Example cortical columns showing target assignment of Ai34-labeled synaptic puncta for Sst^Cre^, Nkx2-1^Cre^, Pv^Cre^ and Nkx2-1^CreERT2^ BC- and ChC-rich regions. (B) Stacked plots depicting interneuron synaptic target preferences for the 4 Cre driver lines in (A). (C) Example images of dissociated cultures in which Sst, BC and ChC are labeled by Ai34 (white) and counterlabeled with target compartments – Kv2 (red), Ankyrin G (green), gephyrin (blue). (D) Stacked plots depicting interneuron target preferences for dissociated cultures using various Cre driver lines (Nkx2-1 for entire MGE lineage, Sst^Cre^ for Sst interneurons, Nkx2-1^CreERT2^ bulk for primarily Pv lineages, BC cultures from Nkx2-1^CreERT2^ BC-rich region, ChC cultures from Nkx2-1^CreERT2^ ChC-rich region. (E) Direct comparison of stacked plots of target preference of cortical sections versus dissociated culture counterpart. (F) Interneuron targeting of soma (red) and dendrite (blue) as a function of cortical depth in cortical columns of Nkx2-1^Cre^, Pv^Cre^, Sst^Cre^, and Sst^Cre^; *Reeler* mice. Grey boxes denote approximate depths of cortical layers. (A) Nkx2-1^Cre^ n=6, 2 cortical columns/animal; Sst^Cre^ n=7, 2 cortical columns/animal; Pv^Cre^ n=8, 2 cortical columns/animal; Nkx2-1^CreERT2^ BC- and ChC-rich n=3, 2 cortical regions/animal. (B) Nkx2-1^Cre^ n=4 cultures, 20-40 tiled regions/culture; Sst^Cre^ n=4 cultures, 20-40 tiled regions/culture; Nkx2-1^CreERT2^ bulk culture n=3, 20-35 tiled regions/culture; Nkx2-1^CreERT2^ BC-rich n=4; Nkx2-1^CreERT2^ ChC-rich cultures n=4. (A, B) 2-way ANOVA, post hoc Tukey’s test. * p<0.05, ** p<0.01, ***p<0.001, ****p<0.0001. (F) Nkx2-1^Cre^ n=6; Pv^Cre^ n=8; Sst^Cre^ n=7; Sst^Cre^; RL/RL n=5.

Second, we assessed targeting in dissociated cultures from Nkx2-1^Cre^, Sst^Cre^ and Nkx2-1^CreERT2^ BC-enriched and ChC-enriched regions (Figure 3C,D). Here, we found a trend towards more dendrite targeting between Sst^Cre^ and Nkx2-1^Cre^ cultures and a significant increase in soma targeting between Nkx2-1^Cre^ and Nkx2-1^CreERT2^ cultures, consistent with known target preferences for interneuron subclass labeled in each line. A direct comparison between Sst, BC and ChC cultures showed that the target preferences between each subclass was significantly different from one another.

Next, we compared targeting in cortical sections versus their counterpart in culture (Figure 3E). With Sst^Cre^, overall targeting was similar between cortical cultures and cortical sections suggesting that Sst interneuron targeting relies mainly on intrinsic mechanisms. With cultured BCs target fidelity was not as robust in cultures compared with cortical sections and we observed a trend towards decreased in soma targeting compared to sections. Similarly, AIS targeting was decreased in ChC cultures.

We then examined targeting in cortical sections as a function of cortical depth (Figure 3F). We found that soma and dendrite target is not even across layers. As expected, we observed a strong enrichment of dendrite targeting in layer 1. This was present in both Nkx2-1^Cre^ and Sst^Cre^ driver lines and reflects the fact that both label the Sst lineage that innervates pyramidal dendritic tufts. With Pv^Cre^, we found an increase in soma targeting in layers 2/3 and a decrease in soma targeting in layer 6. This was also observed in the Nkx2-1^Cre^ driver line but not as pronounced since the Cre driver labels both Pv and Sst lineages. The enrichment of soma targeting was absent in Sst^Cre^. It seemed likely that layer variations in interneuron synaptic targeting reflected differences in the underlying postsynaptic environment. To test this, we crossed Sst^Cre^; Ai34 to the *Reeler* mutant allele. In *Reeler* mutants, cortical structure is severely disrupted with deficits in inside-out radial migration and pyramidal cell polarity ^29^. This was confirmed by immunohistochemistry, which revealed a general inversion in cortical layering and disrupted pyramidal cell polarity (Supplemental Figure 9). Analysis of synaptic targeting as a function of cortical depth in Sst^Cre^; RL/RL mice revealed gross differences in targeting across layers, including a loss of dendrite-exclusive targeting in superficial cortex and a loss of structure along the superficial to deep axis (Figure 3F).

### Unsupervised Analysis Reveals Interneuron Synaptic Subgroups

In order to assign a target for each synapse, ilastik takes into consideration measurements of the syn-tdT puncta itself and its spatial and intensity relationship to the 4 raw confocal channels and the 11 probability channels that define pre- and post-synaptic compartments. In total, ilastik assigns 696 metrics to each syn-tdT object, generating a highly multidimensional dataset from the millions of synapses analyzed. Importantly these synaptic features were not pre-selected based on their ability to discriminate the three targeting classes from each and were therefore suitable for unsupervised analysis.

To begin, we performed an unsupervised analysis on synapses from Nkx2-1^Cre^ and Sst^Cre^ dissociated culture datasets. Synapses in culture are effectively a single cell layer and as a result more uniform in their immunolabeling, making the dataset inherently less noisy. Second, we reasoned that downstream analysis would be easier in culture since the relationships between synapses and targets would be easier to interpret in monolayer culture rather than the more complex environment of a cortical section.

We used a standard workflow for unsupervised analysis, which consisted of dimensionality reduction followed by clustering. Briefly, we took the object feature metrics computed by ilastik as input, z-scored each feature, performed dimensionality reduction using a standard autoencoder, and ran the Louvain community detection algorithm on the Euclidean distances in autoencoder space to cluster objects into putative classes of boutons (more details in Materials and Methods). To test the robustness of the autoencoder, we calculated the reconstruction loss using mean squared error and observed similar losses across multiple seeds using different machines (3.7% ± 0.12 across 40 seeds). Having established that the autoencoder approach was robust for dimensionality reduction, we then generated Uniform Manifold Approximation and Projection (UMAP) plots. Overlaying the supervised classification predictions on the UMAP plot revealed that the different target classes (dendrite, soma and AIS) group together, indicating that the 2-D embedding for visualization can distinguish between syn-tdT puncta with different targets (Figure 4A). For our bouton clustering, we used two assessments to optimize over parameters: 1) comparison of intercluster and intracluster dispersion maximizing the Calinski-Harabasz score and minimizing the Davies-Bouldin score, and 2) segregation of annotated target classes, with each individual cluster enriched for a single target (Supplemental Figure 10).

**Figure 4.**
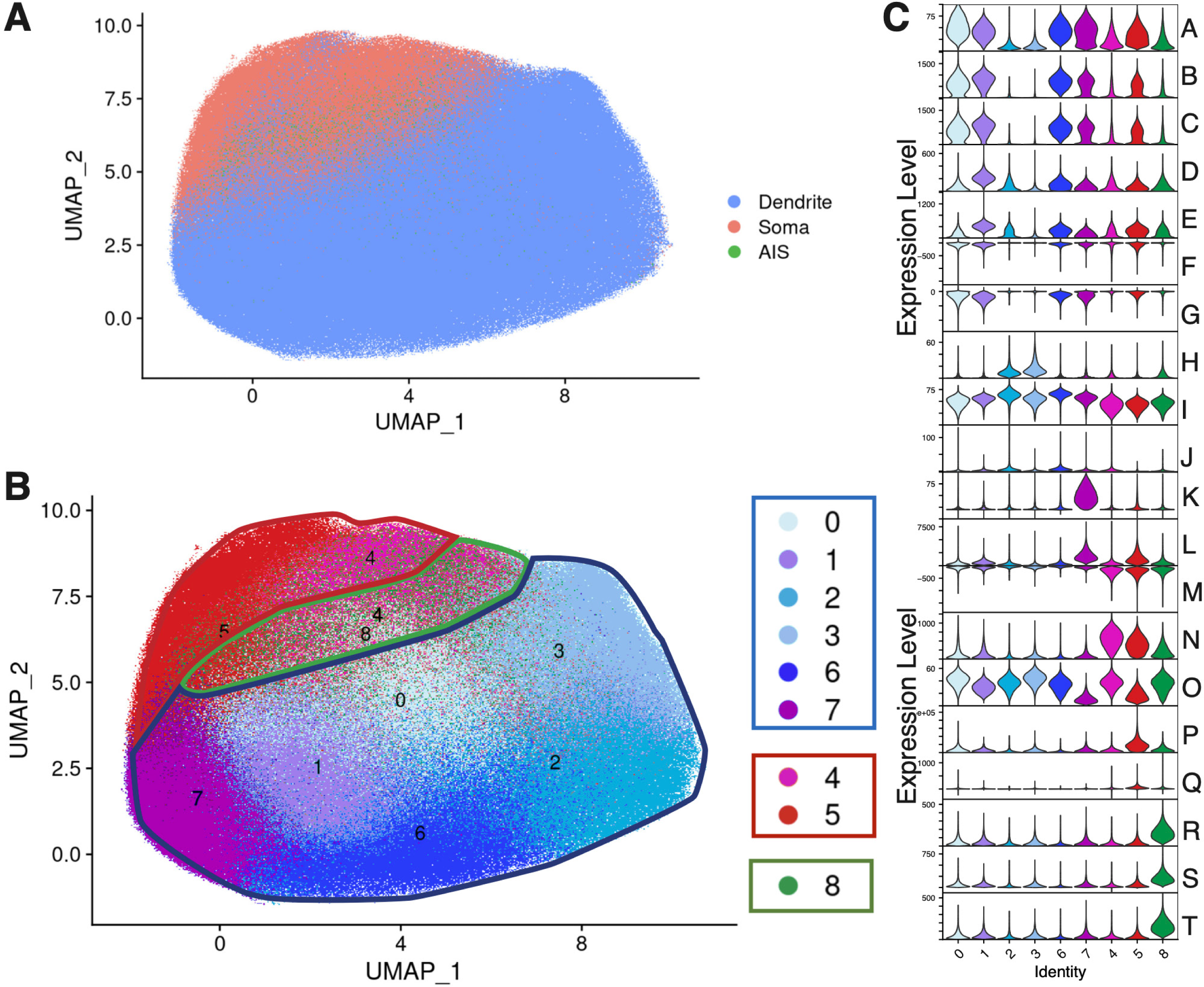
Unsupervised analysis of Ilastik-derived synaptic features reveals the presence of synaptic subgroups. (A) UMAP visualization of synapses from the in vitro culture data set, generated from autoencoder-based intermediate layer nodes as the reduced dimensional space. Points are colored by Ilastik-derived (supervised learning) target calls. Co-segregation of soma-, dendrite- and AIS-targeting synapses is apparent in this visual representation. (B) Same plot as in A, but with points colored by cluster identity following Louvain community detection to identify clusters, showing subgroups enriched in dendrite- and soma-targeting synapses. (C) Violin plots showing the distribution of ilastik-derived feature values, with features selected based on maximal discriminability between each cluster and all the others shown in (B). Features are ordered to highlight that clusters are distinguished by combinations of features with continuous values rather than sharply-defined breaks. Feature Legend: A) mean intensity of gephyrin probability; B) variance of intensity in gephyrin probability; C) Covariance of channel intensity of gephyrin probability; D) variance of intensity in neurite probability; E) Variance of intensity in bouton probability; F) covariance of channel intensity soma probability versus gephyrin probability; G) covariance of channel intensity gephyrin probability versus non-AIS Ankyrin G probability; H) minimum intensity in non-AIS Ankyrin G probability; I) mean intensity in postsynaptic target background; J) kurtosis of intensity in presynaptic background; K) minimum intensity in bouton probability; L) variance of intensity in Ai34 raw channel; M) covariance of channel intensity in Kv2 versus presynaptic background probability; N) variance of intensity in soma probability; O) mean intensity in neurite probability; P) total intensity in RFP soma probability; Q) covariance of channel intensity in Kv2 raw versus RFP soma probability; R) variance of intensity in AIS probability; S) covariance of channel intensity in Ankryin G raw versus AIS probability; T) covariance of channel intensity in AIS probability.

Through this process, we arrived at our static model grouping synapses in dissociated culture, in which we identified 9 clusters: 6 dendrite-targeting subgroups, 2 soma-targeting subgroups, and 1 AIS group (Figure 4B). Based on these subgroups, we identified the image-based features that most clearly delineated each subgroup (Figure 4C) and the top 20 features with the greatest variation between subgroups (Supplemental Figure 11).

### Spatial Distribution of Synaptic Subgroups

Although we used multiple metrics and some prior information to ensure that our bouton clusters were robust, the question remained as to whether these groupings have biological significance beyond the major targeting classes. To tackle this question, we examined the spatial organization of our subclusters, with the goal of assessing whether there was non-random distribution. This was possible by tracking boutons from each synaptic subgroups back onto their source images via their x, y, z coordinates within the source tile.

We first ran a spatial compactness analysis, whereby we assessed the extent to which boutons from a subcluster were more likely to be spatially proximal to each other, as opposed to boutons from a different cluster. Visual examination revealed that synapses belonging to the same subgroup were likely localized in space. We then performed a systematic compactness analysis, whereby we identified the 10 nearest neighbors for each bouton (using a modified Louvain approach), and assessed whether these neighbors were enriched for boutons of the same cluster, as compared to the null distribution of all bouton clusters (Figure 5A). By organizing the mean local enrichment scores for each bouton class versus all others, we made 2 observations: First, the spatial distribution of synaptic subgroups is significantly non-random, in that boutons of the same cluster tend to group together in space. This is not surprising for the AIS cluster, given the spatial compartments its boutons target as compared to the other clusters. By contrast, it was not a given that the two soma-targeting clusters and the dendrite-targeting clusters would also show spatial segregation with respect to their counterparts. Second, it was clear that some subgroups were spatially correlated or spatially anti-correlated with other subgroups (Supplemental Figure 12A), suggesting that individual clusters might show higher-order spatial organization.

**Figure 5.**
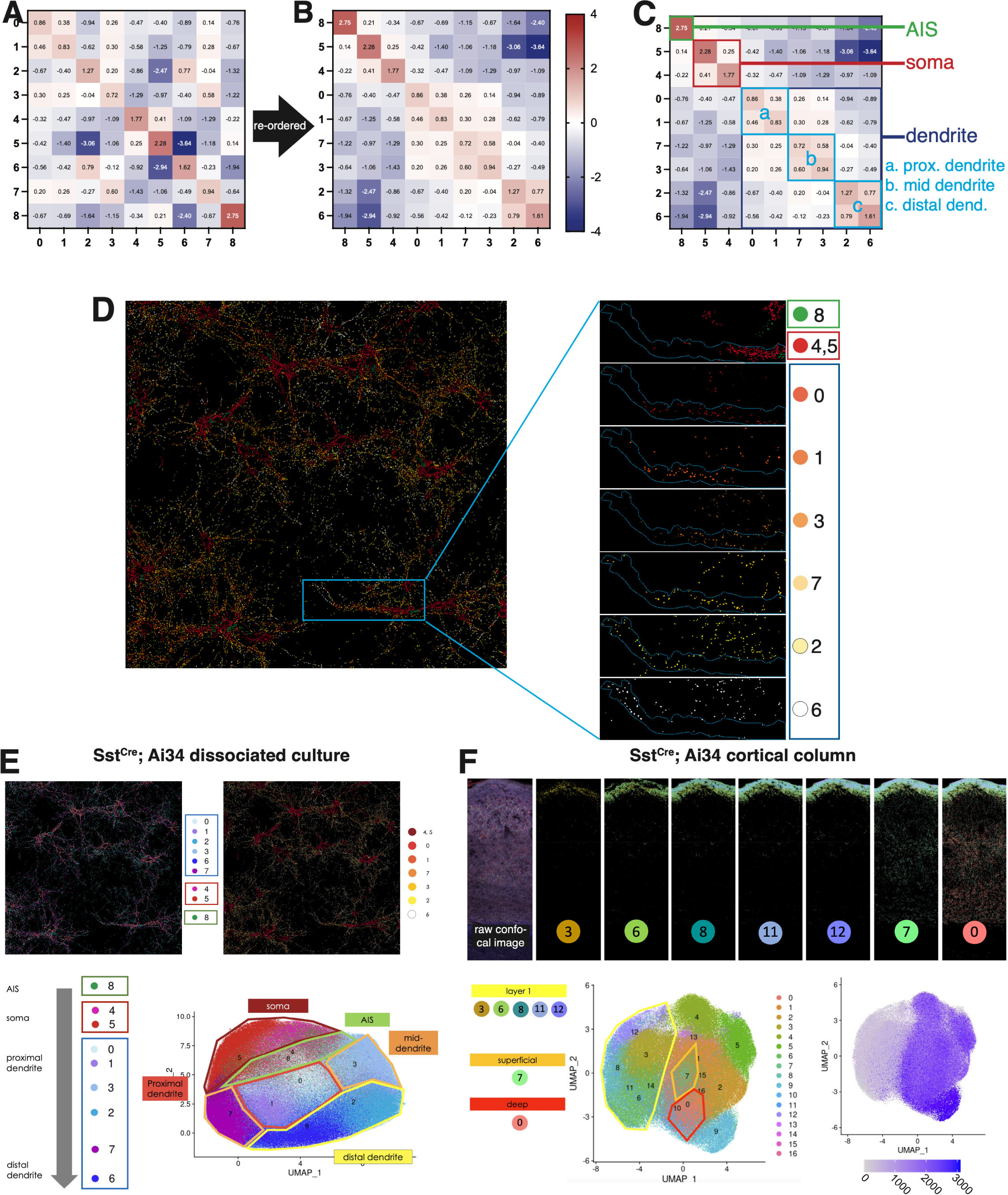
Analysis of synaptic subgroup spatial compactness reveals sequential spatial relationship between dendrite-targeting subgroups along proximal-to-distal axis of target dendrite. (A, B) Modified Louvain analysis of x, y, z coordinates of synapses belong to each cluster (0-8) divided by random distribution and plotted as a enrichment matrix. (A) Enrichment matrix showing spatial relationship when clusters are ordered numerically from 0 to 8. (B) Re-ordered enrichment matrix showing spatial relationship when clusters re-order to maximize off-diagonal sum values. (C) Re-ordered enrichment matrix with cell compartments (AIS, soma, proximal, mid and distal dendrite) denoted on plot. (D) Soma- and dendrite-targeting subgroups re-colored in red-orange-white heatmap. Boxed region rendered sequentially from proximal to distal dendrite (top to bottom). Dotted blue line outlines bundle of dendrites originating from somata on right of image. (E) Synaptic subgroups rendered in original colors and red-orange-white heatmap onto Sst^Cre^ dissociated culture. Spatially-segregated dendrite-targeting subgroups indicated on UMAP of synaptic clusters. (F) Synaptic subgroups rendered onto Sst^Cre^ cortical column. Left, spatially-segregated dendrite-targeting subgroups indicated on UMAP of synaptic clusters. Observation confirmed by (right) UMAP overlaid with cortical depth of synapses in pixels. Analysis of ∼1.45 million synapses obtained from Nkx2-1^Cre^ n=4 and Sst^Cre^ n=3 cultures. Analysis of 2 Sst^Cre^; Ai34 cortical columns consisting of ∼20 million synapses.

Given that these spatial relationships are founded on subcellular targeting of distinct compartments, we next asked whether there is a natural, optimized sequence for these bouton classes. The values at the super- and sub-diagonals designate the spatial relationship between one subgroup and the next on the matrix. We reasoned that maximizing the sum of all super- and sub-diagonals across the matrix would result in a sequence where adjacent subgroups are organized to maximize this spatial correlation (Supplemental Figure 12). We calculated the sums super- and sub-diagonals for every permutation of the 9 synaptic subgroups to empirically determine the optimal sequence. This analysis yielded an optimized sequence of 8, 4, 5, 0, 1, 3, 7, 2, 6 (Figure 5B, Supplemental Figure 12B). Re-ordering the enrichment matrix according to this optimized sequence shows AIS cluster followed by the 2 soma-targeting subgroups, then the 6 dendrite-targeting subgroups (Figure 5C). A closer examination of the dendrite subgroups revealed additional substructure. We found 3 pairs of subgroups that showed high spatial correlation with each other and, with each successive pair, an increasingly greater anti-correlation with soma and AIS subgroups. That is, our analysis indicated that dendritic subgroups are spatially arranged along the proximal to distal axis of the dendrite (in relation to the soma). Indeed, when we re-color the dendritic subgroups as red-orange-white heatmap and re-map them onto the raw image, the proximal-to-distal arrangement is clear (Figure 5D).

Our findings suggest that arrangement of synaptic subgroups reflects a topographic map of the postsynaptic environment. To further test the robustness of this finding, we performed an independent clustering of the boutons, using a different dimensionality reduction technique (principal component analysis, PCA) and a different clustering approach (k-means clustering on Euclidean distance in PC space). The clusters we identified using this alternative approach were largely similar to those obtained from the autoencoder+Louvain approach, and when the optimal consensus sequence for PCA+K-means was mapped onto raw images, we observed similar soma-enriched and dendrite-enriched clusters along the proximal to distal axis (Supplemental Figure 13). Satisfied with the robustness of our clustering approach, we next tested whether dendrite-targeting subgroups were similarly arranged on the dendrites of pyramidal neurons in cortical sections.

In the cortex, pyramidal neurons are highly polarized. They bear an apical dendrite that projects superficially, often ramifying in layer 1, and basal dendrites that are found in deeper layers. We performed a similar synaptic cluster analysis on a Sst^Cre^; Ai34 cortical column (Figure 5F). We noted that there were approximately twice as many synaptic subgroups identified compared to cultures. We mapped the synaptic subgroups back onto the column and found 5 dendrite-targeting subgroups that were highly enriched in layer 1, corresponding to the distal dendrite of pyramidal neurons. We also found subgroups that were enriched in superficial and deep layers, which may correspond to proximal and mid dendrite. Therefore, the dendritic subgroups we identified in culture and those identified in cortical sections both localize to dendrites in a proximal-to-distal manner. Taken together, our analysis revealed parallels in dendrite-targeting subgroups along the proximal-to-distal axis in dissociated culture and in cortical sections (Figure 5E, F).

### Soma-Targeting Synaptic Subgroups

Next, we examined the 2 soma-targeting subgroups (4 and 5) (Figure 4B). We first tested if they localized to different portions of the soma in the way that dendrite subgroups localized along the dendrite (Supplemental Figure 14). We mapped subgroups 4 and 5 back onto raw images and found that 4 and 5 clustered apart from one another. Next, we examined a synaptic metric that is a proxy for proximity to AIS and found no differences between subgroups 4 and 5. Taken together, it suggests that subgroup 4 and 5 do not map to different subdomains of the soma. We next examined the spatial distribution of subgroups 4 and 5 in relation to the raw confocal channel for Kv2, which labels soma and proximal dendrite. This revealed that subgroups 4 and 5 contact different postsynaptic soma (Figure 6A). We quantified this feature by soma and found the distribution to be tri-modal (Figure 6B): soma that contained >70% subgroup 4 (type 4 soma), soma that contained >70% subgroup 5 (type 5 soma), and soma that contained ∼50% mixture of subgroups 4 and 5 (type 4/5 soma). scRNA-seq studies have identified multiple subtypes of BCs and it was possible that different BC subtypes might form synapses that correspond to soma subgroups 4 and 5 or might specifically contact soma type 4, 4/5 and 5. To test this, we cultured Ai34-labeled BCs at low density with unlabeled cortical neurons, which allowed us to examine the axonal arbors of individual BCs. In all individual BCs examined (n=15), we found both subgroup 4 and 5 synapses were formed by the same cell and that the same BC formed synapses onto soma that were type 4, 4/5, and 5 (Supplemental Figure 15).

**Figure 6.**
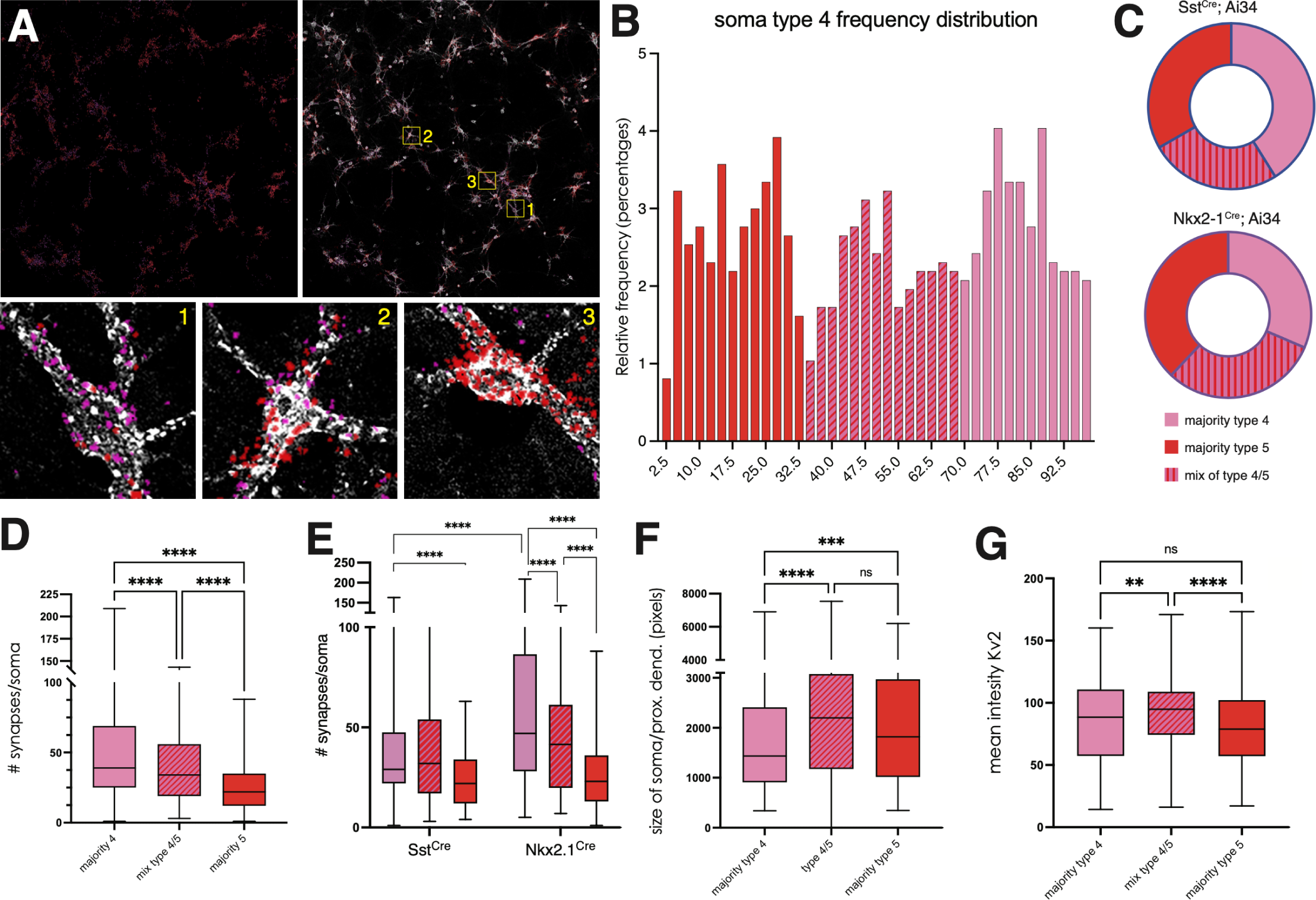
Soma-targeting synaptic subgroups label distinct subpopulations of postsynaptic neurons. (A) Soma targeting synaptic subgroups 4 (pink) and 5 (red) rendered onto original image without (left) and with (right) Kv2 labeling of soma and proximal dendrite (white). Boxed regions show postsynaptic cells that predominantly receive subgroup 4 synapses (type 4 cells), receive a mixture of subgroups 4 and 5 synapses (type 4/5 cells) and predominantly subgroup 5 synapses (type 5 cells). (B) Frequency histogram of proportions of synaptic subgroup 4 and 5 innervation showing the delineation or type 4, type 4/5 and type 5 postsynaptic cells. (C) Proportion of type 4, 4/5 and 5 cells found in Nkx2-1^Cre^ and Sst^Cre^ cultures. (D-G) Features of type 4, 4/5 and 5 postsynaptic cells. (D) Number of synapses received/soma for each type of postsynaptic cell. (E) Number of synapses received/soma separated by Cre driver. (F) Size of soma and proximal dendrite for each type of postsynaptic cell. (G) Intensity of Kv2 signal for each type of postsynaptic cell. Based on the analysis of 977 somata (Nkx2-1^Cre^ 600 somata from 4 cultures; Sst^Cre^ 377 somata from 3 cultures). One-way ANOVA, posthoc Tukey’s test. * p<0.05, ** p<0.01, ***p<0.001, ****p<0.0001.

This finding suggested to us that types 4, 5, and 4/5 soma represented distinct postsynaptic populations that dictated the nature of soma subgroup synapses that formed onto it. As such, we wondered if Sst interneurons, which also form soma-targeting synapses, but to a lesser degree, also targeted soma in the same fashion. A comparison of Nkx2-1^Cre^ and Sst^Cre^ cultures shows that types 4, 5, and 4/5 soma can be found in the same proportions for both Cre driver lines, consistent with the notion that these are distinct postsynaptic cell types, rather than differences in targeting as a result of interneuron subclass (Figure 6C). If the postsynaptic populations are in fact distinct, we would expect that types 4, 5 and 4/5 cells to exhibit distinct features. To test this, we quantified the number of synapses formed onto types 4, 5 and 4/5 and found that type 4 soma contained significantly more synapses than type 4/5, with type 5 containing the fewest number (Figure 6D). The increase in synapse number for type 4 was only observed in Nkx2-1^Cre^ cultures, not Sst^Cre^ cultures, suggesting that BCs labeled in Nkx2-1^Cre^ cultures were likely responsible for the elevated soma targeting in type 4 (Figure 6E). Next, we examined the size and Kv2 intensity of type 4, 4/5 and 5 soma. We found that type 4/5 was significantly larger and more had more intense Kv2 signal than types 4 and 5. In our dataset, we found a sizeable number of type 4 and type 5 cells that exclusively contained subgroups 4 and 5. We found that these cells contained significantly fewer synapses, had the smallest soma size and had the lowest intensity of Kv2 (Supplemental Figure 16). Taken together, our data suggest that subgroup 4 and 5 soma-targeting distinguishes postsynaptic neuron types (4, 4/5 and 5), which exhibit distinct features and likely represent distinct postsynaptic populations.

### Inhibitory Coverage by Interneuron Subclass

We examined individual Sst, BC and ChCs to test whether interneuron subclasses preferentially employed synaptic subgroups (Supplemental Figure 17). Unsurprisingly, we found that BCs made use of subgroups 4 and 5 (the soma-targeting subgroups) more than Sst and ChC and that ChCs preferentially used subgroup 8 (AIS-targeting). With Sst interneurons, there was a significant increase in the use of subgroup 6, which corresponds to the subgroup that targets the most distal portion of the dendrite. But overall, Ssts, BCs and ChCs in culture make use of all 9 subgroups of synapses to varying degrees. This prompted us to examine the axonal branches of individual interneurons to test whether the sequence of synapses formed along their axons might be distinct across interneuron subclasses.

We mapped subgroups back onto the raw images of individual BCs and Sst and re-colored them as red-orange-white heatmap based on the optimal sequence we determined earlier (Figure 5B, C). We noted that BC axonal branches bore synapse subgroups that were consistent with the optimal synaptic sequence. In contrast, Sst axonal branches bore synaptic subgroups that were often non-sequential (Figure 7A, B). To quantify this, we developed a synaptic ‘sequentiality’ score that accounts for the subgroup identity of each synapse along an axonal branch and counts the average number of ‘jumps’ from the optimal sequence for each successive synapse (Supplemental Figure 18 A-C). For example, if a subgroup 3 synapse is followed by a subgroup 6 synapse, this would constitute a jump of 3 (the optimal sequence is 8, 4/5, 0, 1, 3, 7, 2, 6,) (Figure 7C). We normalized the sequentiality score on a scale of 0 to 1 (1 being highly sequential) and quantified the average normalized sequentiality score for axonal branches from individual BCs and Ssts (n=10 and n=12 cells). We found that BCs axonal branches were significantly more sequential than Sst axonal branches (Figure 7D, E). We tested compared BC and Sst sequentiality to randomized iterations. Both Sst and BC sequentiality are both non-random but BC sequentiality is significantly less random, consistent with our findings (Supplemental Figure 18D, E). A potential confound is that BCs are far more likely to form multiple successive soma-targeting synapses, which would result in successive jumps of 0, artificially inflating sequentiality score. We therefore performed a secondary analysis in which we only scored sequentiality for contiguous stretches of axon containing dendrite-targeting subgroups (Figure 7F). Even with this more stringent measure, BC axonal branches are still significantly more sequential than Sst. We interpreted these data to suggest that BCs form synapses along the processes of postsynaptic partners, thereby resulting in synapses that follow postsynaptic topology and are, thus, more sequential. In contrast, Sst axons form a small number of synapses onto each postsynaptic partner and then ‘skip’ to the next process resulting in synapses that are more randomized.

**Figure 7.**
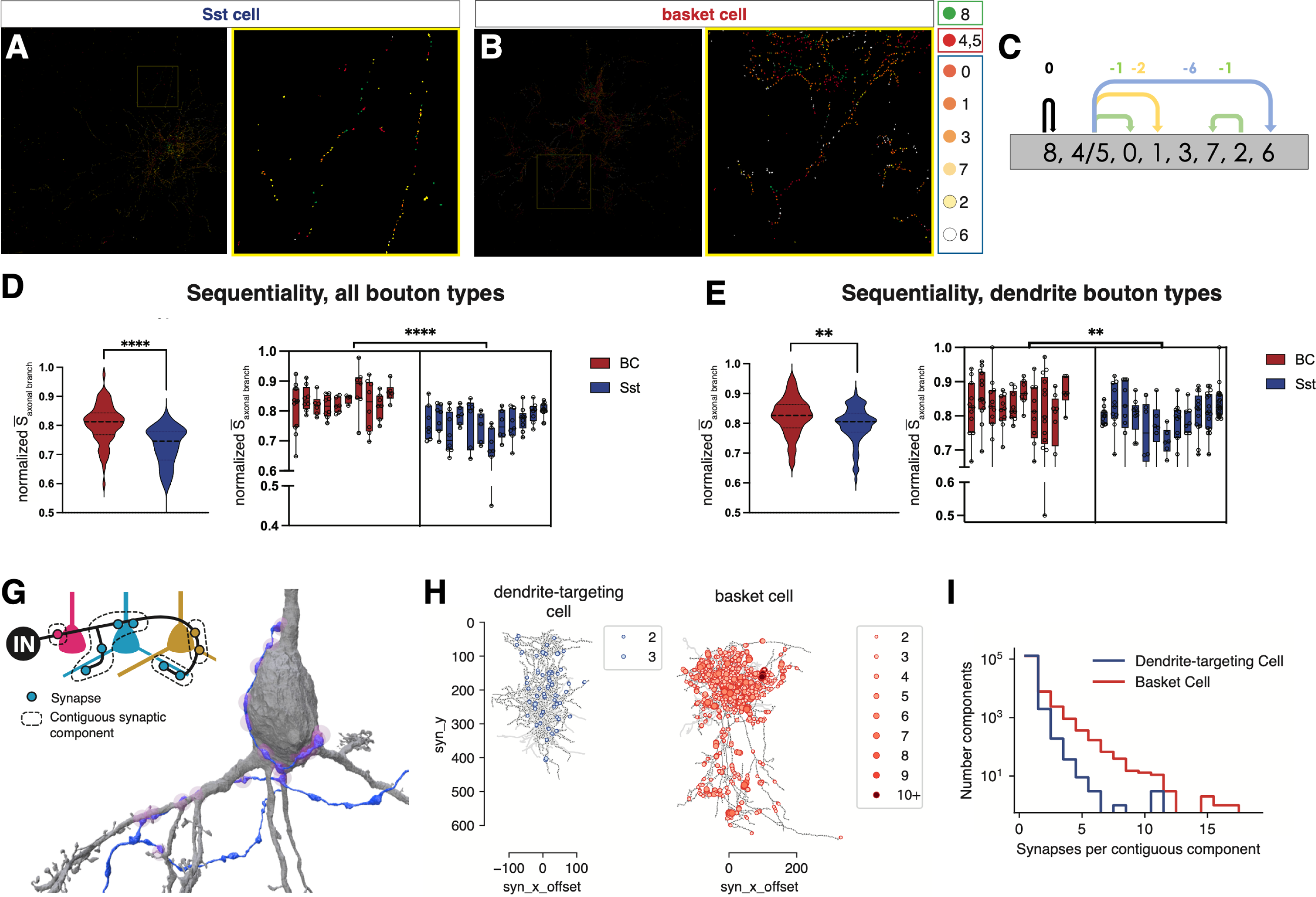
Analysis of Basket cell and somatostatin cell synapses along their axons reveal different strategies for enacting inhibitory coverage. Individually-labeled (B) Sst interneuron and (B) BC with synapse subgroups rendered in red-orange-white heatmap reflecting optimized synaptic subgroup sequence. (C) Calculating sequentiality score from successive synapses along axonal branch by their deviation from optimized sequence (See Supplemental Figure 17 for more details). (D) Frequency (left) distribution of the average sequentiality scores of individual axonal branches from Sst and BCs; 2-way ANOVA p<0.0001. (E) Sequentiality score for all (left; unpaired t-test p,0.0001) and individual cells (right) for BCs (red) and Sst (blue); 2-way ANOVA p<0.0001. (F) Sequentiality score for all (left; unpaired t-test p=0.0016) and individual cells (right) for BCs (red) and Sst (blue) where only contiguous dendrite-targeting subgroups were calculated; 2-way ANOVA p=0.0013. (D-F) Based on the analysis of 10 BCs and 12 Sst individually-labeled neurons with 5-9 axon branches analyzed per cell obtained from 3 BC cultures and 2 Sst cultures. (G) Schematic depicting interneuron (IN) targeting 3 postsynaptic neurons. Dashed lines indicate regions where synapses are formed successively onto the same postsynaptic neuron, which we term contiguous synaptic components. Image on right shows 3D reconstruction of IN (blue) forming contiguous synaptic components (pink) onto pyramidal neuron (gray). (H) Representative wire diagram depictions of axonal trees of dendrite-targeting cell, left (putative Sst) and basket cell (putative Pv). Overlaid with single synapses (gray circles) and contiguous synaptic components ranging from 2-16 synapses (blue or red circles). (I) Frequency distribution of synaptic component sizes for dendrite-targeting (putative Sst, blue) and basket cells (putative Pv, red); Mann-Whitney U test (p<10^-300^).

To test this assertion further, we analyzed single cell reconstructions from a large electron microscopy volume of mouse visual cortex ^30^. A collection of 58 perisomatic targeting cells (putative BCs) and 54 dendrite-targeting cells (putative SSTs) along a column have been extensively proofread ^31^, with hundreds to thousands of synaptic outputs for each cell, focusing only on synaptic outputs targeting excitatory neurons (n=292,673, see Materials and Methods). Note that no chandelier cells were in these reconstructions. To quantify the relationship between synaptic targeting and axonal morphology, for each cell we identified regions of axon in which synaptic outputs target the same postsynaptic cell, which we called contiguous targeting domains (Figure 7G). Each contiguous targeting domain was characterized by its number of synapses (Figure 7H), which ranged from 1–17 synapses. Consistent with our analysis of axon subgroup sequentiality, we found that BCs had significantly larger targeting domains compared to dendrite targeting cells (p<10^-300^, Mann-Whitney U test), owing to a long tail of large components (Figure 7I). Taken together, our analysis of synaptic subgroup sequentiality aligns well with observed connectivity properties observed in the intact cortex.

## Discussion

In this study, we developed an image-based workflow to assign interneuron synaptic targeting at scale and with high concordance with human user scoring. As expected, this approach detected significant differences in targeting across interneuron Cre driver lines both in cortical sections as well as in dissociated culture. This afforded us the opportunity to directly compare target fidelity between cortical interneurons in the intact brain and compare it to the fidelity for interneurons in dissociated culture. Overall, we found that interneuron subclasses in culture showed similar target preference to interneurons in culture but the degree to which targeting was retained varied by subclass: Sst interneurons displayed remarkably similar targeting to Sst interneurons in cortical sections whereas BCs and ChCs in culture had reduced soma and AIS targeting compared to cortical sections, respectively. A recent EM connectomic analysis of interneuron synaptic development found early and persistent dendrite targeting by Sst cells while BC synapses successively refined onto soma over time ^32^. Thus, BCs in culture might represent an arrested state of maturity, which accounts for the decreased soma targeting. Additionally, parvalbumin expression is not robust in vitro (which is why we did not use Pv^Cre^; Ai34 for our culture experiments). Parvalbumin expression is sensitive to neuronal activity levels ^33^ and it is therefore also possible that activity levels in culture may be insufficient or inappropriate for Pv lineage cells to achieve maturation. Similarly, ChCs (the other Pv lineage subclass) also lagged behind their in vivo counterparts in target fidelity. ChCs are thought to have near-absolute target preference for AIS ^13,14,34^, which may suggest that ChCs, like BCs, also fail to fully mature in culture. Indeed, ChC varicosities are pruned and refined over a prolonged period of time ^35^. Nevertheless, we observe clear chandelier cell-like cartridges in culture, which is, to the best of our knowledge, the first described instance of ChC cartridge formation in vitro and suggests that AIS targeting is at least partially under intrinsic regulation. Taken together, it suggests that refinements in culture conditions could help to bring in vitro and in vivo synapse targeting into closer alignment.

Despite limitations, it is notable the extent to which synaptic target fidelity and synaptic features are retained in dissociated culture. We find significant target preferences across Sst, BCs and ChCs in culture and perhaps more remarkably, analysis synaptic subgroups in culture reveals novel interneuron biology that was corroborated in follow up analyses in cortical sections (Figure 5E and F; Figure 7). The parallels between neurons in culture and in the intact brain argue in favor of a prominent role for intrinsically determined cellular programs in the establishment of subdomains in dendrites and interneuron synaptic connections. It also lends credence to the use of in vitro approaches as a simple and tractable model going forwards to address some of the outstanding questions pertaining to interneuron synaptic diversity and specificity.

A key innovation of our approach is in training a model to perform a biologically-relevant computer vision task (assignment of synapses to target compartments), which in turn generated a multidimensional dataset of spatial- and intensity-based synaptic features. This allowed us to assess large numbers of synapses at a population level at the resolution of cellular subcompartments, and to identify novel synaptic subgroups. Coupled with posthoc analysis of the spatial distribution of subgroups, it represents a novel and powerful method to study synapses going forwards, one that can be modified to suit experimental needs.

With this method, we identified dendrite-targeting synaptic subgroups that were organized in a proximal to distal arrangement away from the soma. By performing a similar analysis in cortical sections, we found dendrite-targeting subgroups that specifically mapped onto layer 1 or were enriched in superficial and deep layers, consistent with the notion that a proximal-to-distal arrangement is also present in the intact cortex. Prior work describes the compartmentalized nature of electrical signals in dendrite subdomains ^36–38^. It is thought that these local electrical compartments integrate their signals to encode for complex information ^39,40^. The role of inhibitory input onto these compartments can regulate or even override signals ^41^ and our data suggest that the nature of those inhibitory inputs is distinct across dendritic subcompartments, prompting one to speculate that they might also display different functional properties. Indeed, inhibitory synapses targeting distal dendrite exert a stronger than anticipated inhibitory effect on dendritic activity that is distinct from the effects of proximal dendrite-targeting synapses ^42^.

We also identified further heterogeneity with 2 soma-targeting subgroups. Here, we found that they mapped onto 3 postsynaptic populations that exhibited significant differences in size, Kv2 intensity, and the number of synapses/soma. Taken together, our interpretation is that the postsynaptic neuron plays an instructive role as to the nature of the inhibitory synapse formed onto it - proximal to distal compartments of the dendrite dictate which subgroup of synapse can form while the somata of different neuron populations exhibit a preference for relative proportions of soma-targeting subgroups innervating it. Additionally, we find that Sst, BC and ChC subclasses can form the same subgroups of synapses, albeit at different proportions. Taken together, our findings suggest that interneurons are generalists in the types of synapses they can form, dictated by which compartment or cell soma type they encounter. This is not to say that interneurons play a passive role. To be sure, interneuron subclass (and subtype) identity drives strong target preferences for the subcompartments or the choice of postsynaptic partner ^43,44^. Only once it does form a synapse, our data indicates that the nature of that synapse is determined by the target. Notably, a rich complement of proteins is found in the inhibitory postsynaptic compartment ^45,46^, and therefore the postsynapse is well-positioned to establish a wide selection of synapse types.

In recent years, a number of groups have generated serial EM connectomic databases of mouse neocortex, for example ^31,47,48,49^. Object registration of features belonging to every pre- and post-synaptic neuron and their synaptic connections has offered an unprecedented glimpse into the organization of cortical synaptic connectivity. Our image-based, multidimensional approach serves as a complementary and parallel strategy for studying synapses. Its benefits lie in its broad accessibility (requiring only basic confocal microscopy) and its versatility (compatibility with mouse genetics and immunohistochemistry). These features make it a flexible vehicle for hypothesis testing through mouse genetics (as we did with the *Reeler* mutant), molecular perturbation, or the interrogation of other experimental variables. Findings from our approach can then be cross-referenced with the ground truth provided by EM connectomics. Indeed, this was the strategy we employed to confirm differences in inhibitory coverage between Sst and BCs, which, notably, is also consistent with findings from prior studies ^50,51^.

Our image-based approach also has parallels to previous work that has sought to identify synaptic heterogeneity. One such approach was the use of multiplexed fluorescent labeling of synaptic proteins coupled with super-resolution imaging to assign features to excitatory and inhibitory synapses ^52,53^ and employed unsupervised analysis to identify synaptic subgroups. The strength of this approach lies in the number of synaptic molecules analyzed, which revealed distinct differences in molecular composition across subgroups. In contrast, our study performed 2 stages of analysis – one supervised, the other unsupervised. The supervised analysis was anchored on interneuron synaptic targeting onto postsynaptic compartments, which provided the necessary context for us to analyze the spatial distribution of synaptic subgroups identified through unsupervised analysis. Another study performed a population-level analysis on excitatory synaptic diversity ^54^. They examined the presence and intensity relationships between 2 postsynaptic fluorescently-labeled synaptic proteins, and found synaptic heterogeneity in excitatory neurons throughout the brain. Our study focused on inhibitory synapses in the cortex, investigates their association with cellular compartments, and incorporates both pre- and post-synaptic features in our analysis.

Zhu and colleagues found that excitatory synapse types were spatially enriched across brain structures or within layers of the cortex and hippocampus ^54^. In contrast, we find a different organizing principle: interneuron synaptic subgroups distribute locally in subcellular compartments of the postsynaptic cell and across postsynaptic populations. This contrast makes biological sense given the differences between excitatory and inhibitory synaptic connectivity. Cortical interneurons project locally through repetitive synaptic motifs, so it stands to reason that their complement of synapse types would be patterned locally and reflect local postsynaptic topography (subcompartments of dendrite and different species of postsynaptic soma). Excitatory neurons on the other hand are largely long-range neurons whose efferent synapses project to distal points; Here, synaptic diversity reflecting different functional areas of the brain might be expected. In future, employing a uniform image-based analysis of both inhibitory and excitatory systems could prove highly informative, allowing us to study the spatial distribution of excitatory and inhibitory synaptic subgroups simultaneously and infer how the two systems functionally interact.

There are some caveats that should be considered with our findings. First, our analysis was performed at a population level, which has the downside of blurring individual differences. A good example of this is our use of an optimal sequence of synaptic subgroups, which is very likely a simplified view since the optimal sequence was derived from an overall analysis subgroup spatial distribution, resulting in a ‘best fit’ sequence. In actuality, sequentiality is almost certainly more nuanced, perhaps with parallel sequence types occurring depending on postsynaptic cell type or even across dendritic branches. In our current analysis, we are blind to these types of distinctions and it will require further work in future to carefully resolve. Second, it is possible that interneuron synaptic diversity manifests along multiple dimensions. Our analysis only speaks to the diversity in spatial and intensity relationships between pre- and post-synaptic compartments and there might be parallel or even orthogonal forms of synaptic diversity that we are not detecting in our analysis. By changing the starting parameters and/or inputs of the analysis, different forms of diversity might be uncovered.

## Acknowledgements

We would like to thank Ali Argunsah and Theo Karayannis for their support and guidance during the early stages of the study. Thanks also goes out to Eleftherios Koutsilianos and Anne Cavanagh for their work on related aspects of the experiments. Thanks also to Hynek Wichterle and Franck Polleux for their advice during the preparation of this manuscript. This work was supported by R01NS117695 (EA), R21MH131947 (EA) and the Irma T. Hirschl Foundation (EA).

---

**Supplemental Figure 1.**
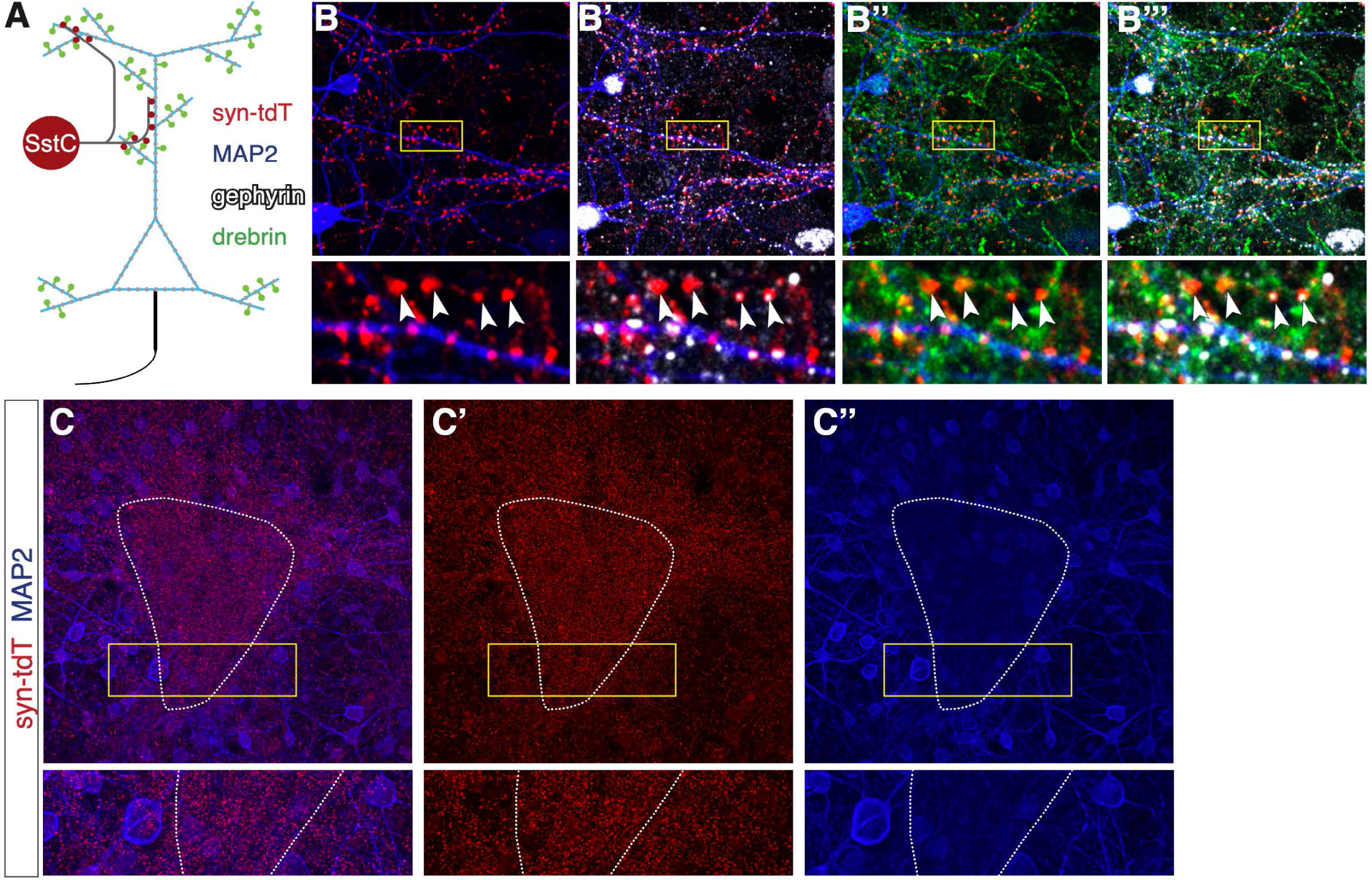
MAP2 is not an ideal postsynaptic marker for dendrite-targeting synapses. (A) Schematic showing labeled compartments in panel B. Synaptophysin-tdT (red), drebrin labeling dendritic spine heads and necks (green), MAP2 labeling spine shafts (blue) and gephyrin (white). (B) Syn-tdT synapses colabeled with MAP2. Arrowheads indicate syn-tdT puncta that are not apposed to MAP2+ shaft. (B’) Syn-tdT, MAP2 and gephyrin shows arrowhead-indicated puncta are apposed to inhibitory postsynapse. (B’’) Syn-tdT, MAP2, drebrin shows arrowhead-indicated syn-tdT puncta are apposed to spin heads and necks labeled by drebrin and are therefore dendrite-targeting synapses. (B’’’) Addition of gephyrin to B’’ shows that drebrin-apposed syn-tdT puncta are also apposed to gephyrin. (C) MAP2 signal is uneven in intensity (area of low intensity circumscribed by dotted line) in areas with high densities of syn-tdT puncta. (C’’) Syn-tdT signal is dense in area of low MAP2. (C’’’) Just MAP2 signal showing uneven signal intensity.

**Supplemental Figure 2.**
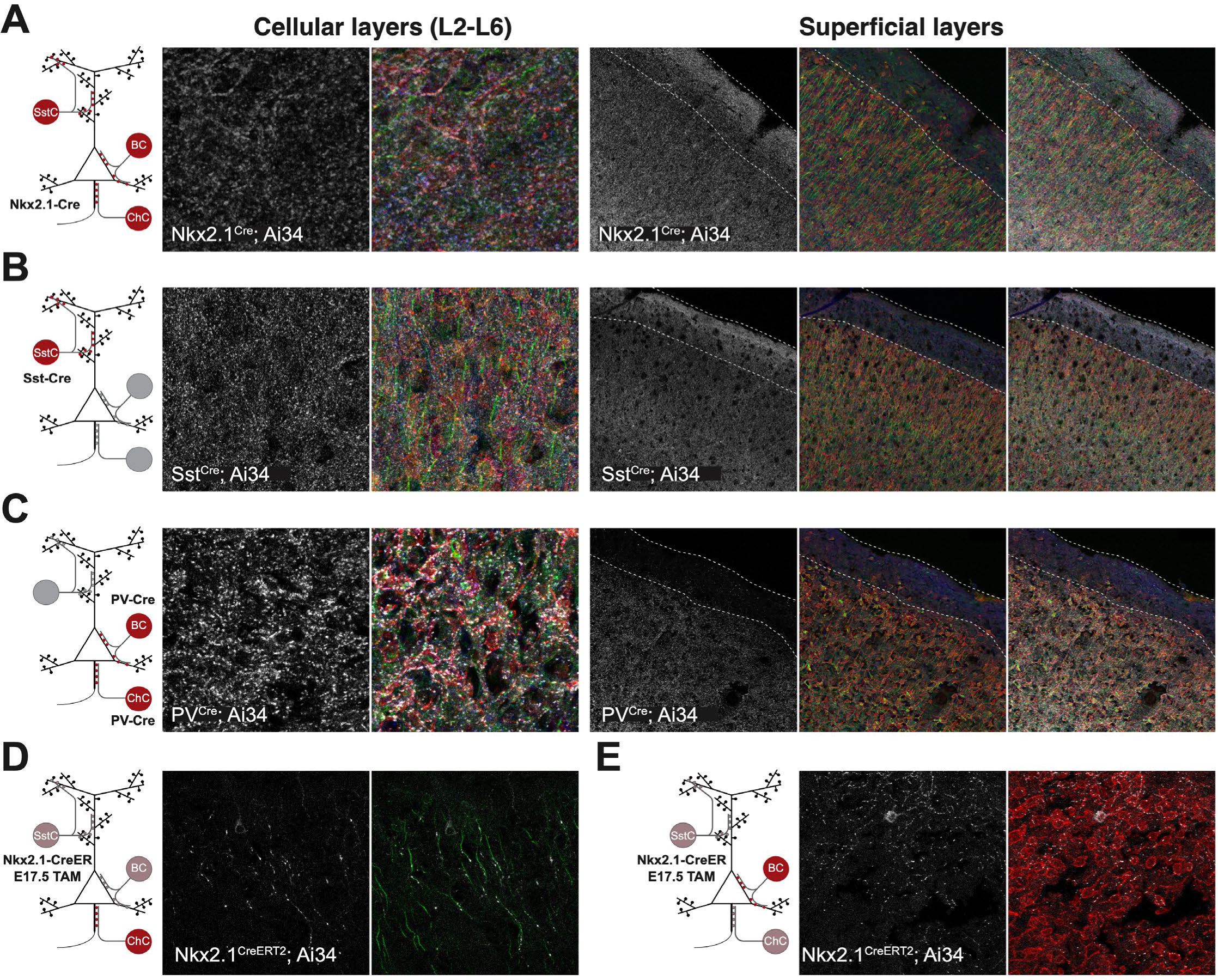
Cre driver lines allow for subclass-specific genetic labeling with Ai34 synaptophysin-tdTomato conditional reporter. (A-C) 3 Cre driver lines (A) Nkx2-1^Cre^, (B) Sst^Cre^, and (C) Pv^Cre^ cortical sections. Left panels show synaptophysin-tdTomato (Ai34) puncta alone and counterlabeled with Kv2 (red), Ankyrin G (green), gephyrin (blue). Right panels show same configuration with layer 1 outlined in dashed white lines. Cortical sections of Nkx2-1^CreERT2^; Ai34 tamoxifen gavage at e17.5 showing synaptophysin-tdTomato puncta alone and counterlabeled with Ankyrin G (green) or Kv2 (red) in regions enriched for (D) chandelier cells and (E) basket cells.

**Supplemental Figure 3.**
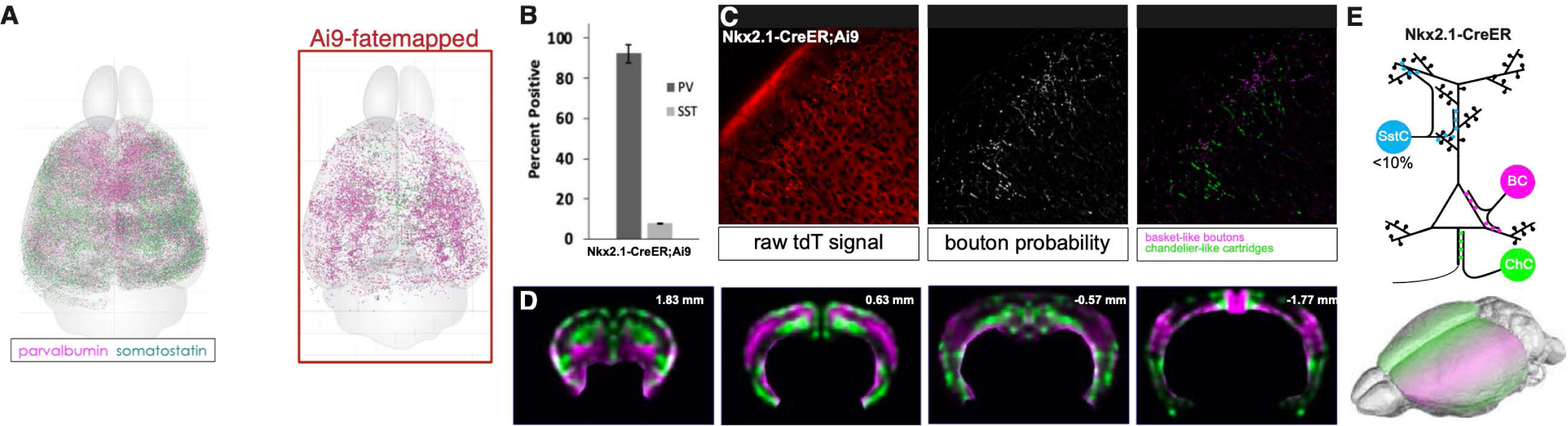
Late embryonic tamoxifen gavage of Nkx^CreERT^^2^; Ai34 mice labels Pv lineage interneuron neurons with spatial segregation between chandelier cells and basket cells. (A) Coronal cortical sections from Nkx2-1^CreERT2^; Ai9 e17.5 tamoxifen mice (n=8) immunolabeled for Pv and Sst. Coronal sections registered to Allen CCF and rendered as 3D model. Left panels shows all Pv (magenta) and Sst (green) cells identified. Right panel shows only tdTomato+ cells. (B) Quantification of tdTomato+ Pv and Sst cells for all sections analyzed. (C) (left) Sample field of tdtomato+ Ai9 signal in coronal section before and after (middle) ilastik processing to generate probability map for synaptic puncta and ImageJ macros to (right) distinguish between chandelier cell cartridge puncta (green) from remaining puncta (magenta). (D) Heatmap composite from 8 mice of regions enriched for chandelier cartridge puncta (green) and BC puncta (magenta). Cortex on the right shows approximate regions enriched for ChCs (green) and BCs (magenta), which informed our dissections for dissociated cultures and imaged regions for target assessment in subsequent experiments.

**Supplemental Figure 4.**
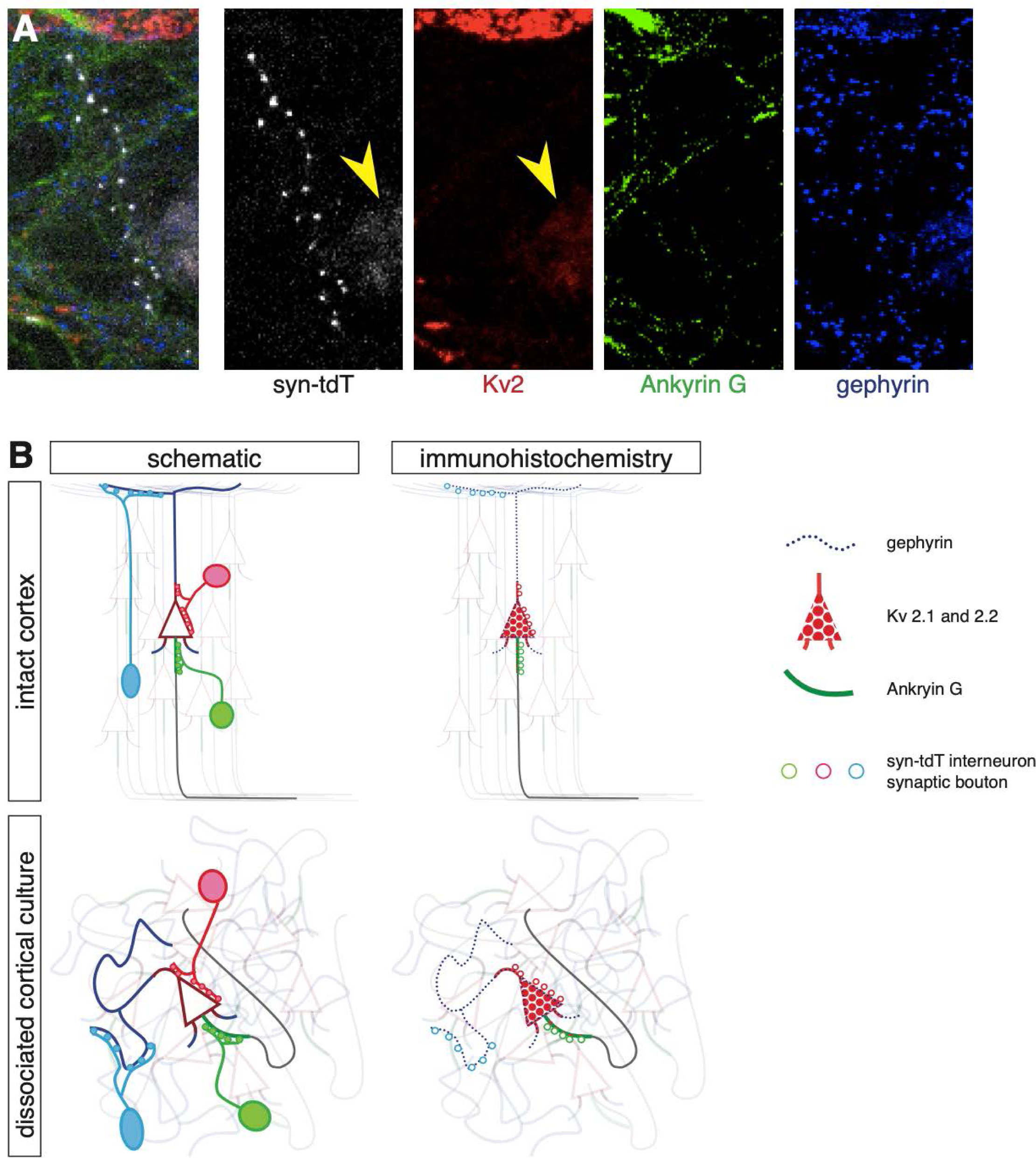
Immunohistochemistry for pre- and post-synaptic features is an imperfect representation of features needed for image-based model analysis. (A) 4 channel confocal image of dissociated Sst^Cre^; Ai34 cortical culture – synaptophysin tdTomato (white), Kv2 (red), Ankyrin G (green), gephyrin (blue). Yellow arrowhead in syn-tdT panel indicates tdTomato signal in soma Sst interneuron, with corresponding Kv2 signal (also yellow arrowhead). Kv2 signal is punctate and does not completely fill in soma and proximal dendrite. Ankyrin G signal is varied: high levels in AIS, lower levels in neurite processes. (B) Schematic showing ideal target compartments vs. immunolabeled compartments – more details in Supplemental Figure 5.

**Supplemental Figure 5.**
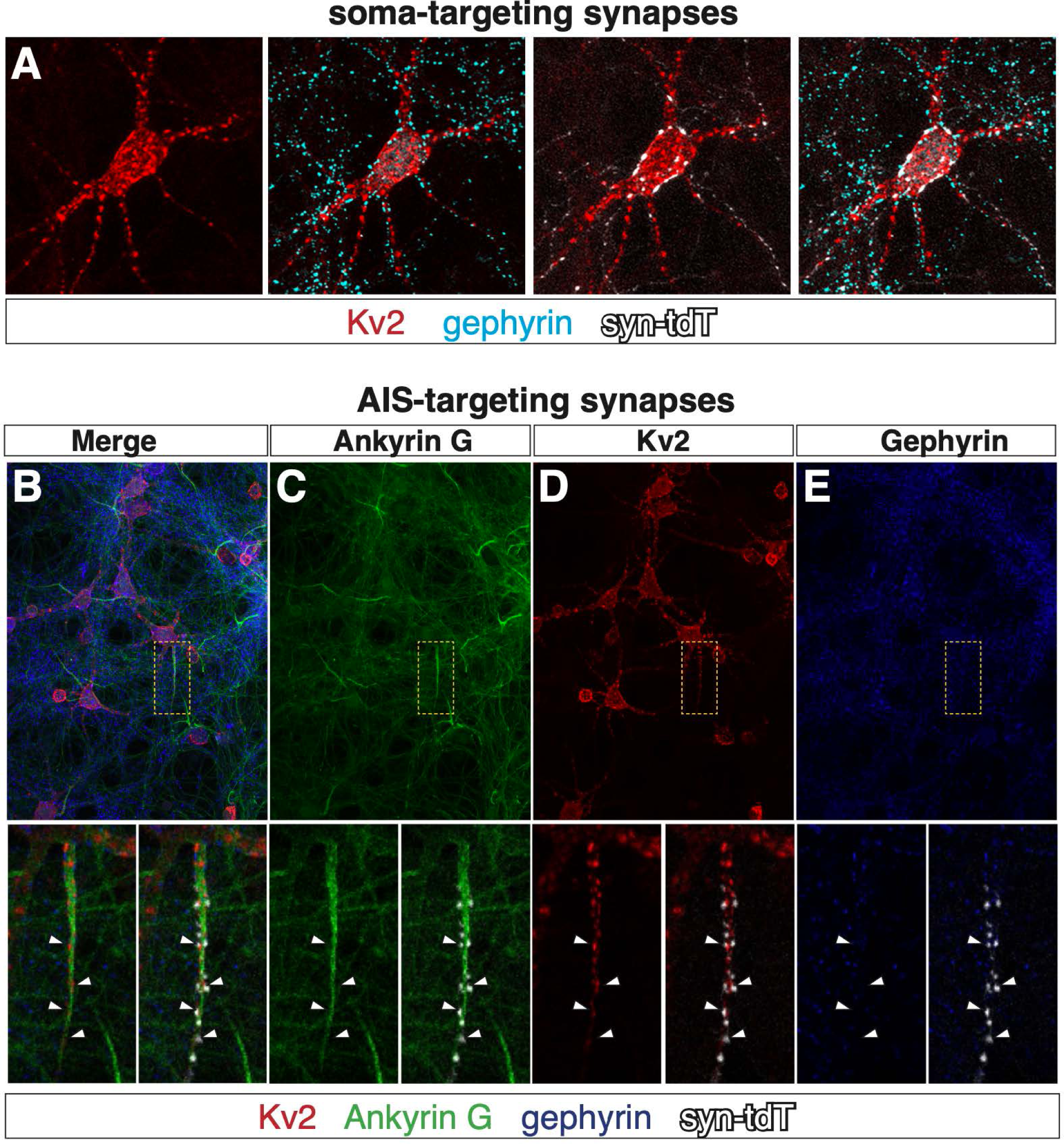
Immunolabeled target compartments for soma/proximal dendrite and axon initial segment are defined by context and multiple channels. (A, B) Immunolabeled dissociated culture neurons - Kv2 (red), AnkyrinG (green), gephyrin (blue) and syn-tdT puncta (white). (A) General immunolabeling pattern for soma and proximal dendrite target compartment. Note punctate Kv2 signal (red), and presence of gephyrin puncta that define target compartment and syn-TdT puncta innervation of soma and proximal dendrite (white). (B) General staining pattern for axon initial segment, merged image of Kv2, Ankyrin G, gephyrin and syn-tdT puncta. (C) Ankyrin G signal at the AIS. Ankyrin G is a canonical marker of the AIS. Note that this antibody also stains neurites in general at a lower intensity, and is excluded from Kv2-rich regions of the AIS. (D) Kv2 signal at AIS. Note that Kv2 localizes to the AIS, where it is found in apposition to AIS-targeting syn-tdT puncta (white). (E) Lack of gephyrin immunolabel at AIS. Gephyrin clustering is a canonical marker of GABAergic post-synaptic densities (PSDs). The gephyrin antibody we use in this study (mAb7a from Synaptic Systems) recognizes gephyrin that is phosphorylated at S270. Gephyrin localized to PSDs at the AIS is not phosphorylated at S270, as indicated by the lack of signal in apposition to syn-tdT puncta.

**Supplemental Figure 6.**
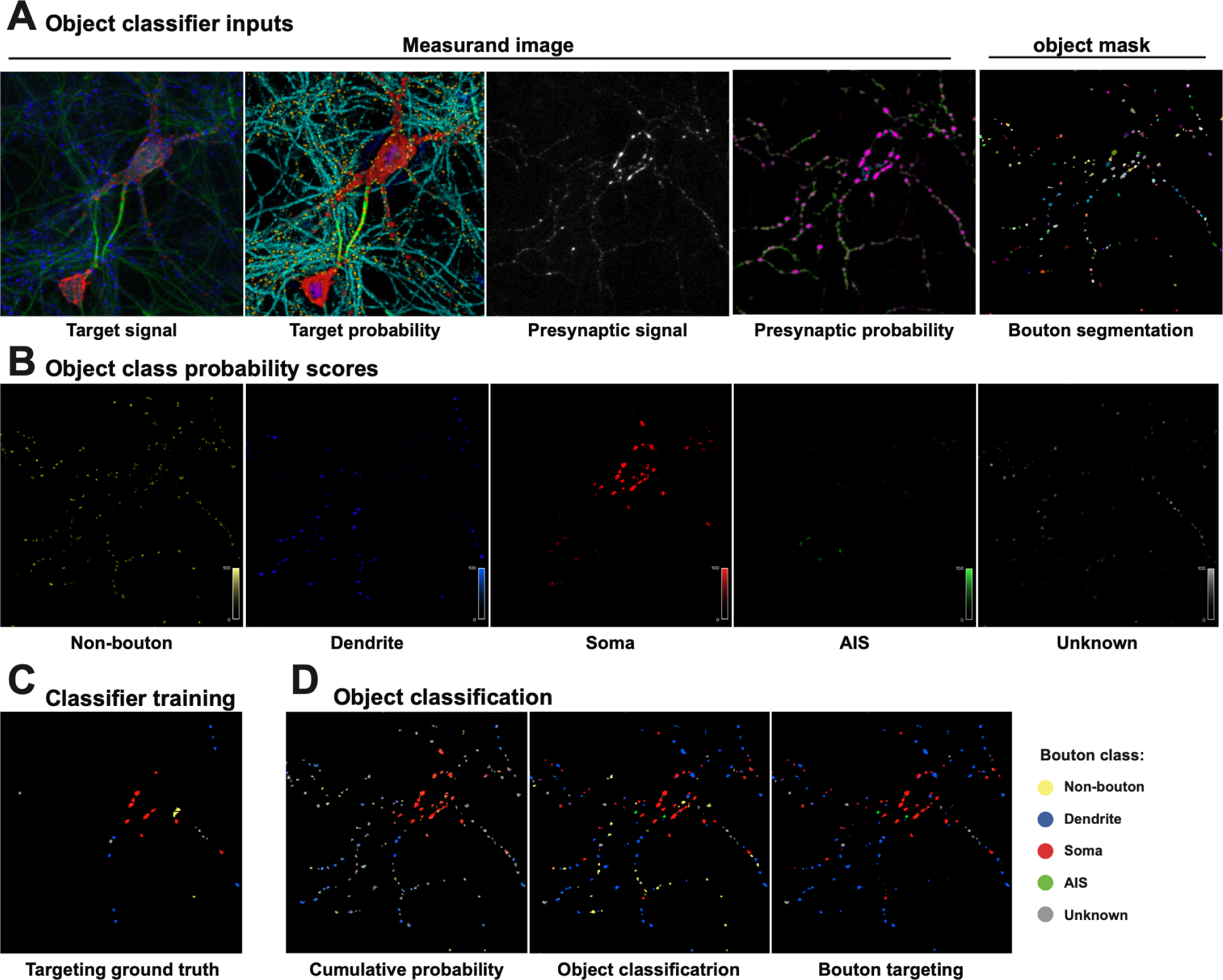
Target assignment for segmented syn-tdT puncta objects. (A) Target assignment (object classification) draws from inputs consisting of the raw and probability mapped target channels (collectively called the measurement basis image) and the object mask (segmented syn-tdT puncta). (B) Target assignment (Object probability score) for the five object classes: non-bouton, dendrite-, soma-, AIS-, and unknown-targeting. Object training of syn-tdT puncta objects. (C) user-trained ground truth data points. (D) (left) Heatmap depicting target probability for all 5 targets, (middle) assigned target based on probability score, (right) finalized assigned targets (non-bouton objects removed).

**Supplemental Figure 7.**
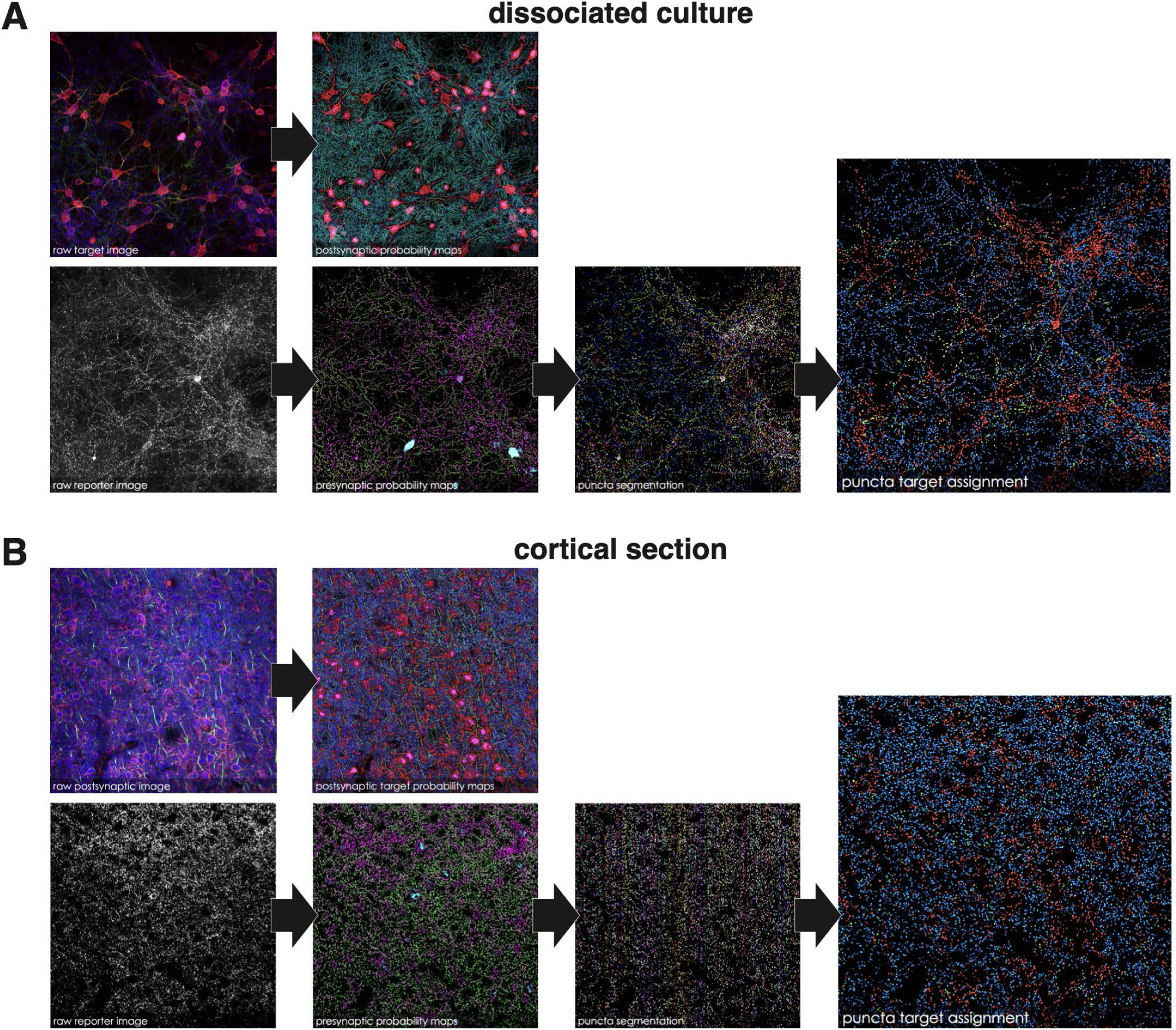
Representative performance of image-based model for synaptic target assignment in dissociated cortical cultures and in cortical. sections. Confocal tiles of (A) dissociated cortical culture and (B) cortical sections processed through various stages of image-based model for assigning synaptic targets. Left, raw confocal channels converted into 7 postsynaptic (top) and 4 presynaptic (bottom) probability channels. Presynaptic channels used for object segmentation of syn-tdT+ puncta. Right, syn-tdT objects assigned targets (red=soma/proximal dendrite; green=AIS; blue=dendrite).

**Supplemental Figure 8.**
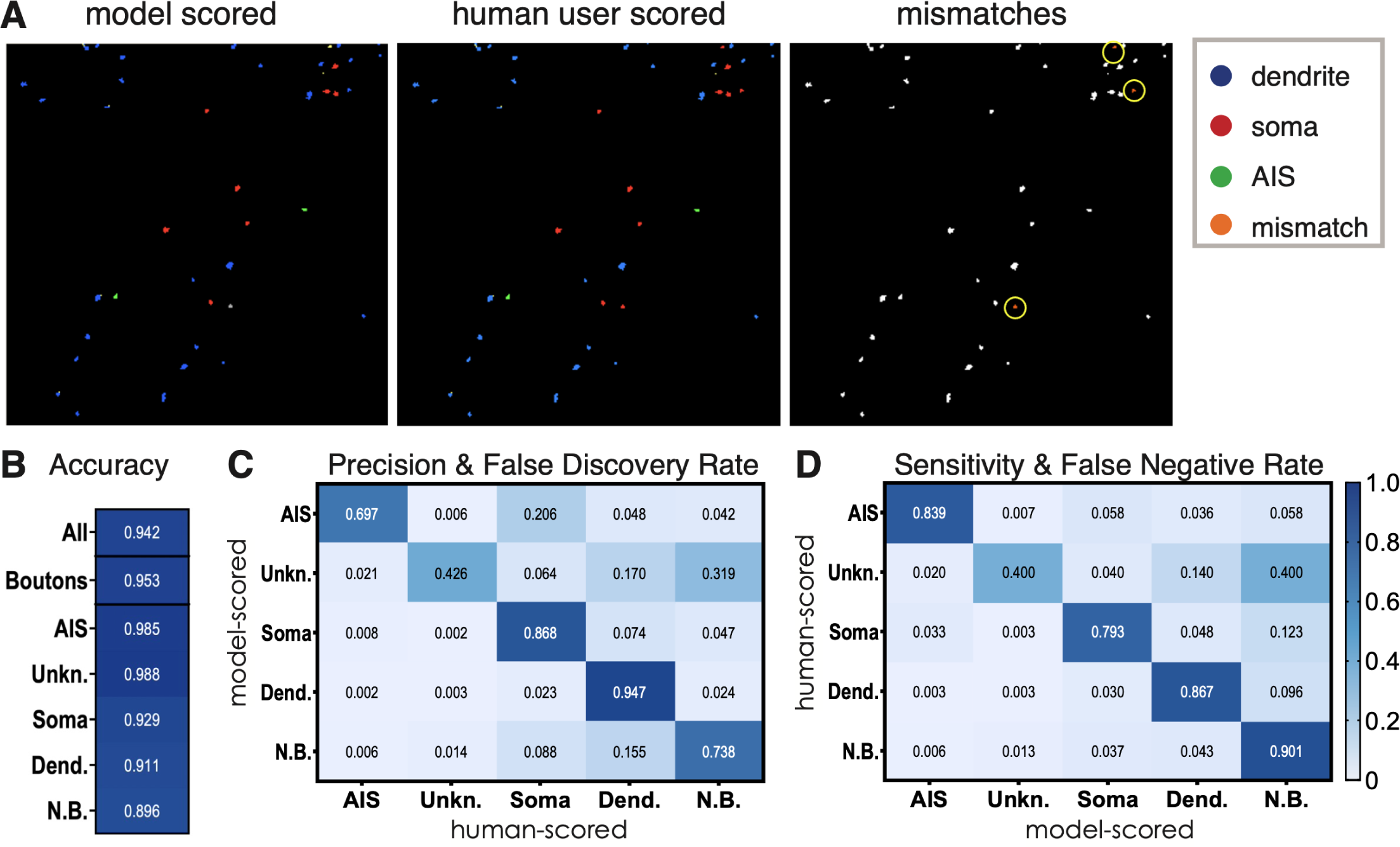
Comparison of model performance versus user assignment of synaptic targets shows strong concordance. (A) Example field of randomly-selected syn-tdT puncta used for comparison between human user and model following target assignment (red=soma/proximal dendrite; green=AIS; blue=dendrite). Orange puncta circled in yellow indicate mis-matches between model and user target assignment. (B) Accuracy of model compared with expert human user scoring. Accuracy is percentage of concordance between human user and model calls. Categories listed in column: all objects, all objects that are boutons, AIS, unknown targeting, soma, dendrite and non-bouton. (C) False discovery rate (off diagonals) shows rates of non-concordance across categories when the model correctly assessed that the object was a positive call but its call did not concord with human user. Precision is represented by the values along the diagonal, which is a measure of the model’s ability to correctly call true positives. (D) False negative rate (off diagonals) shows the rates of non-concordance between model and user when the model incorrectly assesses that the object is a positive call. Values on the diagonal represent the sensitivity, which is a measure of the model’s ability to detect true positives.

**Supplemental Figure 9.**
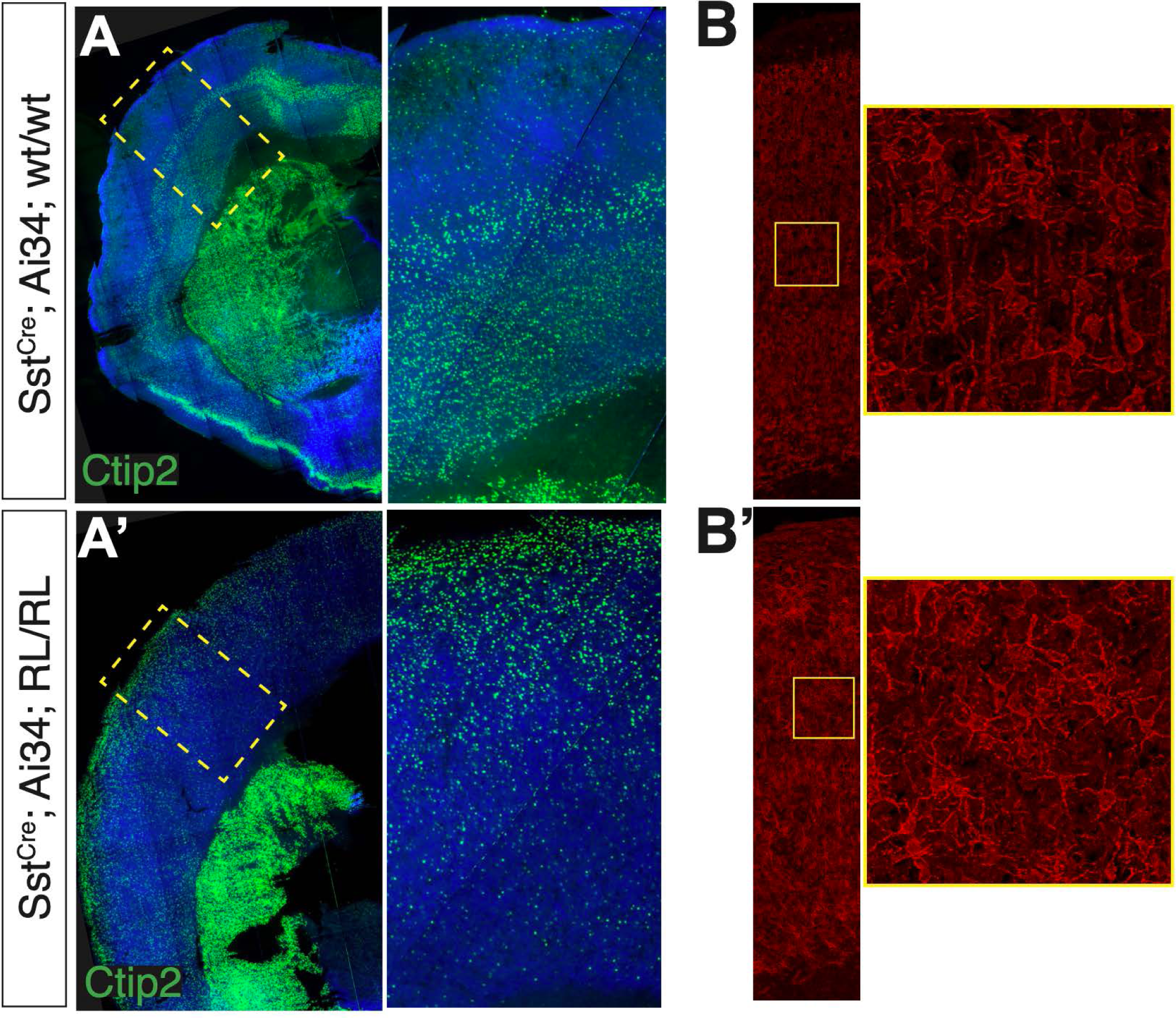
Reeler mutant cortex exhibits inverted pyramidal cell layering, defects in polarity, and randomization of synaptic compartmental target across cortical layers. Coronal cortical sections of P28 Sst^Cre^ mice that are (A) wildtype and (A’) *Reeler* homozygous mutants immunolabled for layer 5-enriched pyramidal cell marker, Ctip2 (green) shows disordered and largely inverted cortex. Kv2 immunolabeling of (B) wildtype and (B’) *Reeler* homozygous mutants shows disruptions in pyramidal cell polarity in mutants.

**Supplemental Figure 10.**
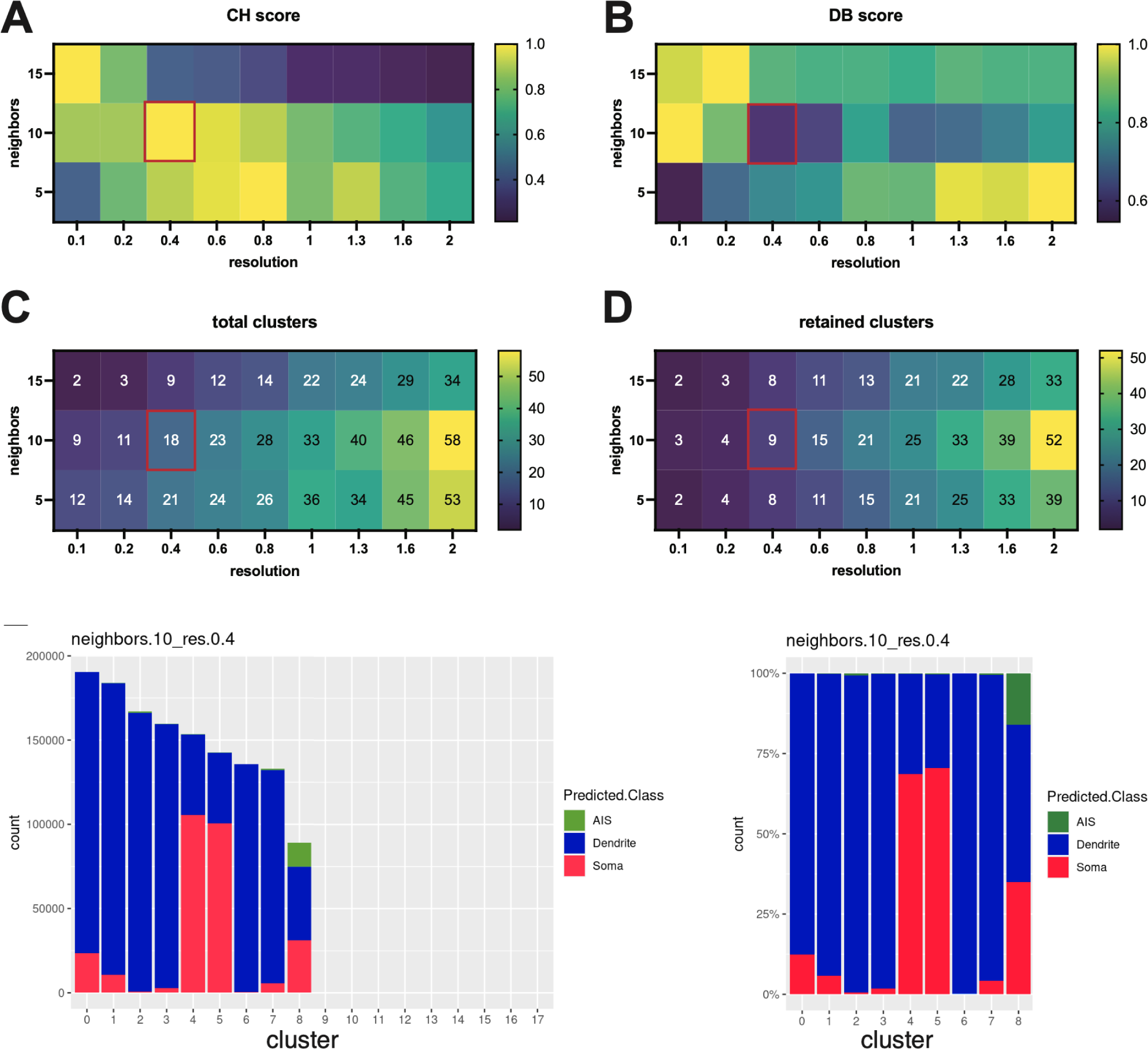
Empirical determination of K and resolution results in the identification of 9 synaptic subgroups. (A) Computed Calinski-Harabasz (CH) score as a function of varying k neighbor value and resolution. (B) Computed Davies-Bouldin (DB) score as a function of varying k neighbor value and resolution. (A, B) To determine optimal k and resolution, we determined where the local maximum for DB coincided with the local minimum for CH – K=10, resolution = 0.4. (C) Computed of number clusters resulting from vary k neighbor values as a function of resolution. (D) Computed number of clusters where the synaptic puncta within the cluster constituted a minimum of 0.5% of the total number of puncta (termed ‘retained clusters, a cut-off for noise). (E) Frequency stacked plot showing the relative abundance of each cluster and their target assignment composition. Note that past cluster 8 cluster abundance is negligible. (F) Target assignment score stacked plot set to 100%.

**Supplemental Figure 11.**
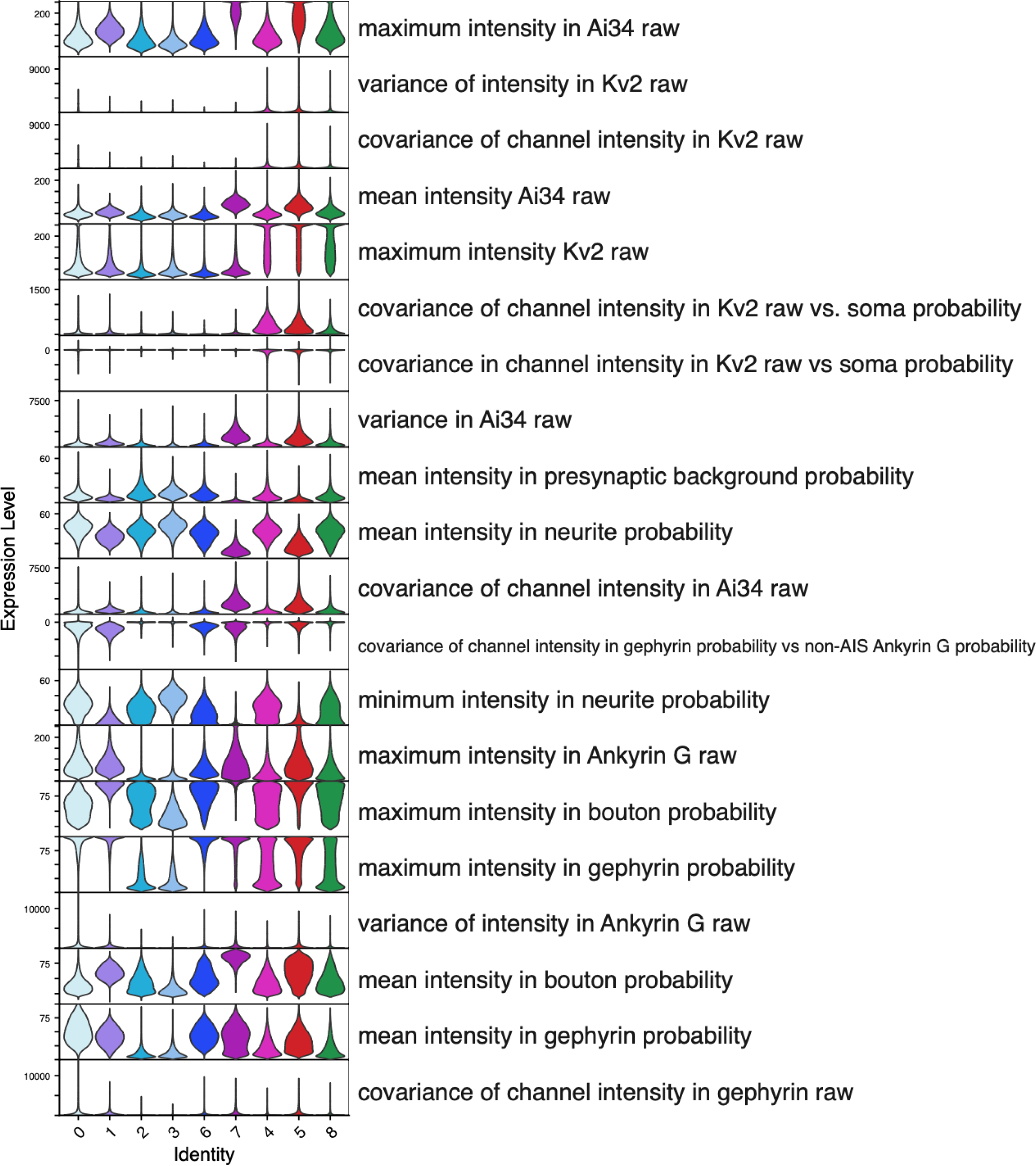
Top-20 highest variance ilastik-derived synaptic features across all 9 identified subgroups.

**Supplemental Figure 12.**
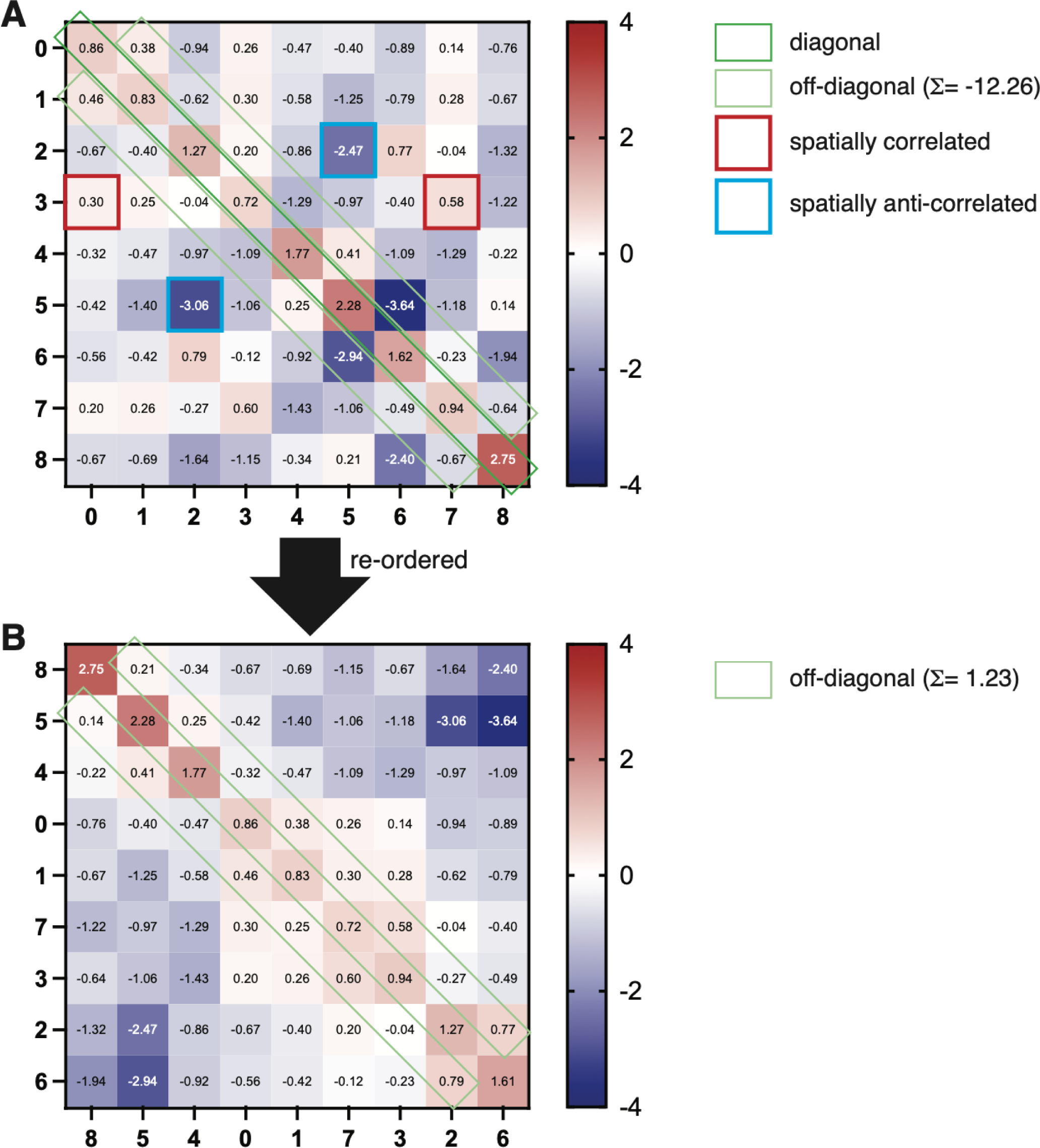
Spatial correlation (spatial compactness) enrichment matrix re-ordering process based on maximal sum of off-diagonal values. (A) Modified Louvain analysis (10 nearest neighbors divided by nearest neighbors assuming random distribution) plotted as an enrichment matrix. Values greater than 0 represent spatial correlation above random distribution and values less than 0 represent spatial anti-correlation vs. random. The matrix displays a *directional* relationship and therefore must be read in a specific manner, namely row to column. Starting with row *i* and reading the value in column *j* asks how many of class *i*’s k-nearest neighbors are coming from class *j*. Doing the inverse, i.e., starting from column *j* and reading the value in row *i*, does *not* provide any information about the inverse relationship. Rather, the inverse relationship is depicted in the cell at column *i*, row *j*. That is, the diagonal values (dark green box) represent the spatial compactness within a cluster whereas the off-diagonal values (light green box) represent the spatial compactness across adjacent clusters. Additionally, cluster compactness across non-adjacent clusters can be found cross-referencing column and row, which shows that some clusters are spatially-correlated (outlined in red box) or spatially anti-correlated (outlined in blue box). We hypothesized that there is an optimal spatial sequence across all the clusters and therefore empirically determined the (B) optimal cluster sequence by calculating the sum of the off-diagonal values for all cluster permutations. In (A), the off-diagonal sum for clusters arranged in numerical order is −12.26. (B) In the optimized cluster sequence, the off-diagonal sum is 1.23.

**Supplemental Figure 13.**
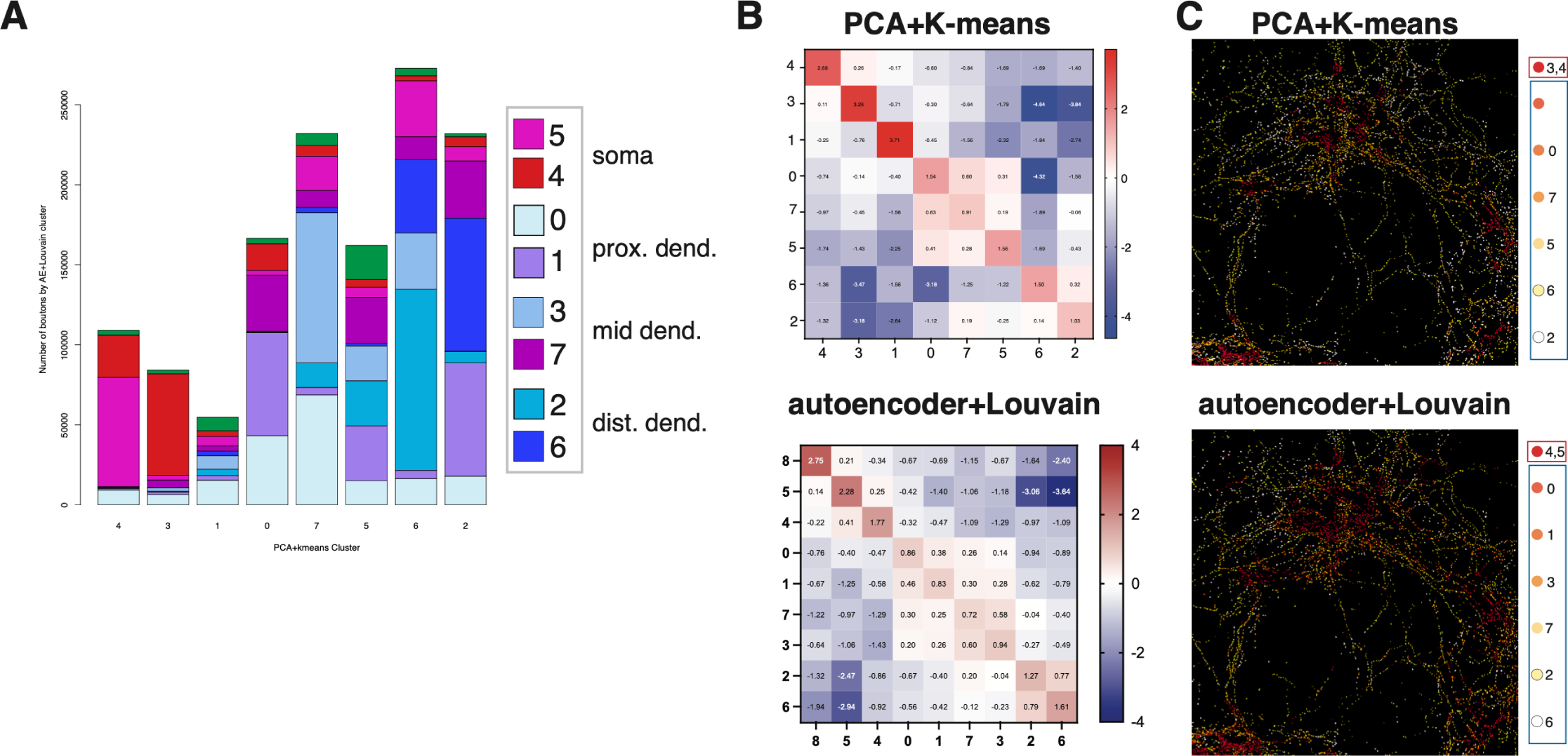
Assessing the robustness and reproducibility of unsupervised analysis of synaptic feature dataset. (A) Stacked bar plot composition of autoencoder+Louvain derived cluster composition for each PCA+K-means derived cluster. Note the general enrichment of soma, proximal, mid and distal dendrite autoencoder+Louvain clusters distributed onto PCA+K-means clusters with similar spatial segregation (B) Enrichment matrix of spatial compactness scores arranged according to empirically-determined optimized sequence for PCA+K-means clusters (top) and autoencoder+Louvain clusters (bottom). (C) Representative Nkx2-1^Cre^; Ai34 culture overlaid with PCA+K-means clusters (top) and autoencoder+Louvain clusters (bottom) colored as a red-orange-white heatmap according to optimized sequence in (B).

**Supplemental Figure 14.**
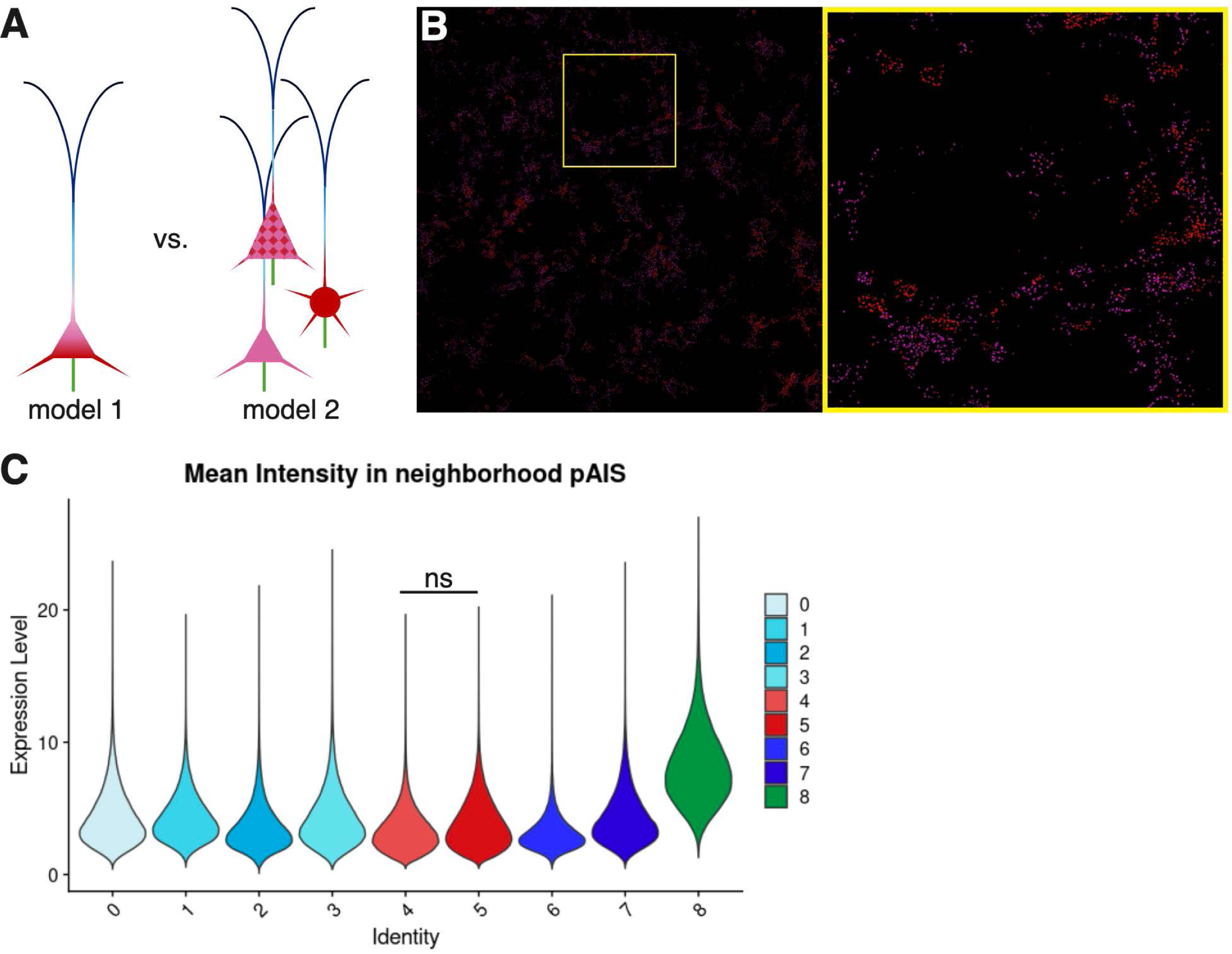
Spatial arrangement of soma-targeting synaptic subgroups. (A) Schematic of 2 competing models for how soma-targeting subgroups might be spatially arranged on target soma. The model on the right depicts subgroup 4 and 5 synapses arranged on somata in a graded fashion with enrichment across different subcompartments of the soma. The model on the left depicts differential enrichment of subgroup 4 and 5 across different populations of postsynaptic neurons. (B) Subgroups 4 (pink) and 5 (dark pink) mapped back onto original image without Kv2 signal. Arrangement of subgroups 4 and 5 is clumped, supporting the model on the left in (A). (C) ilastik-derived metric showing spatial correlation between each synaptic subgroup and the probability map for AIS. No significant difference (1-way ANOVA) between subgroups 4 and 5 suggest that mapping of 4 and 5 onto individual soma is not spatially enriched across soma compartments.

**Supplemental Figure 15.**
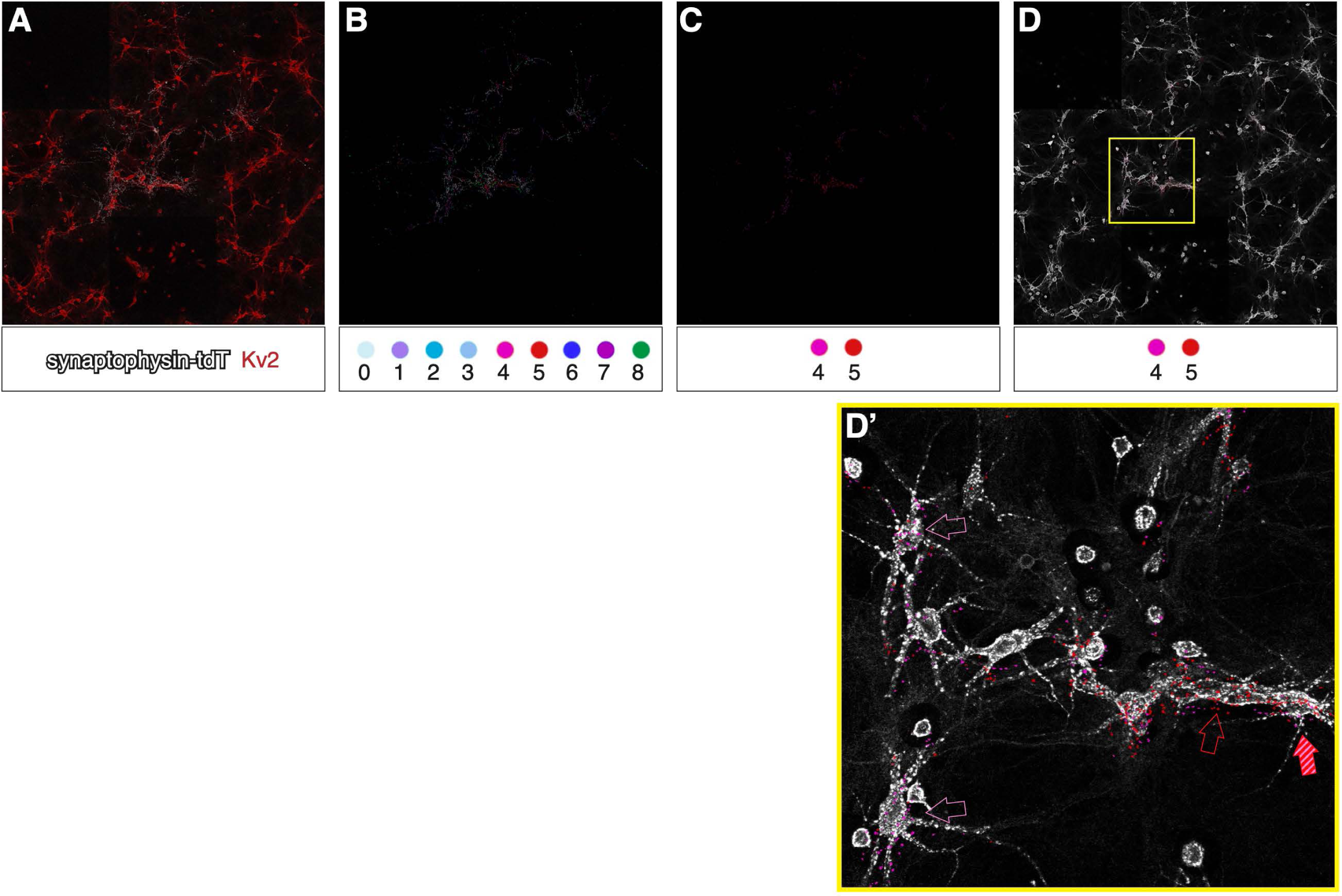
Individually-labeled basket cells form synapses onto types 4, 4/5 and 5 somata. (A) Individually-labeled BC cultured from BC-rich region of Nkx2-1^CreERT2^; Ai34 mice – Syn-tdT puncta (white), Kv2 (red). (B) All synapse subgroups mapped onto the same individually-labeled BC. Just subgroups 4 and 5 mapped on (C) without and (D) with counterlabel Kv2 (white). (D’) Enlarged region of box in (D) shows that the same BC forms synapses onto type 4, type 4/5 and type 5 somata.

**Supplemental Figure 16.**
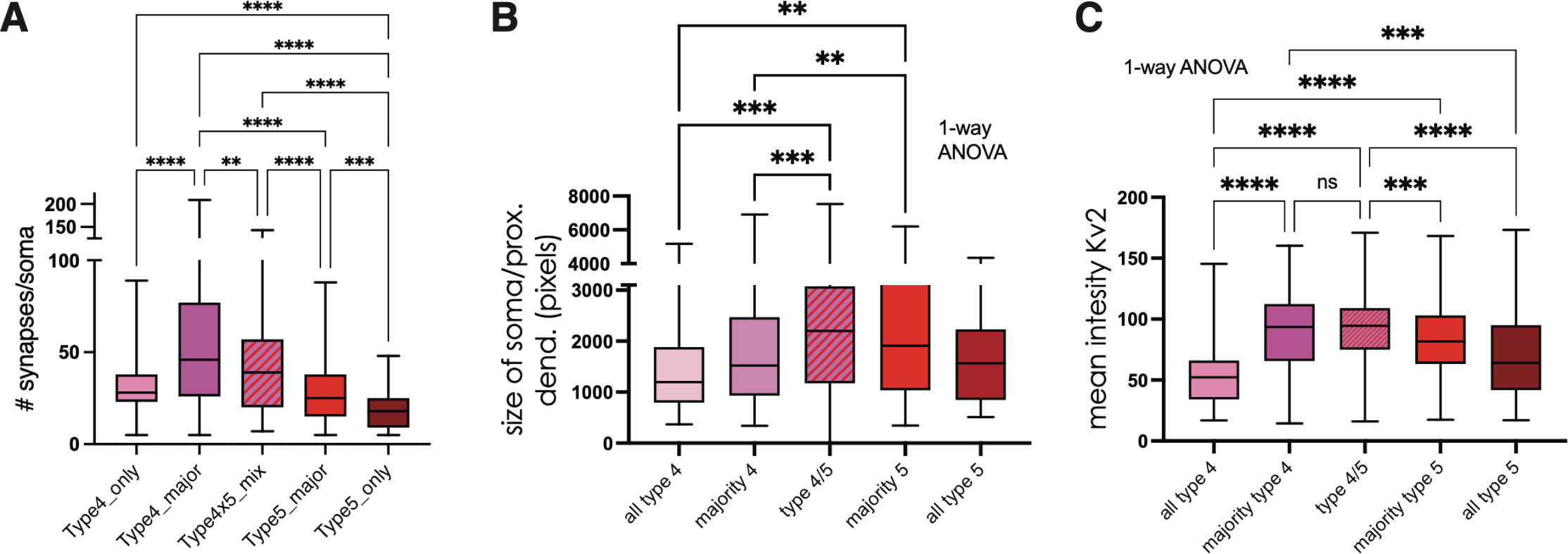
Features of types 4, 4/5 and 5 postsynaptic cells are distinct. A sizeable fraction of type 4 and type 5 postsynaptic cells contain exclusively subgroup 4 and 5 synapses, respectively. These cells are named type 4 only and type 5 only. (A-C) Analysis of features of type 4 only, mainly type 4, type 4/5, mainly type 5 and type 5 only cells for the following measurements: (A) # synapses/soma; (B) size of soma and proximal dendrite; (C) intensity of Kv2 signal. Based on the analysis of 977 somata (Nkx2-1^Cre^ 600 somata from 4 cultures; Sst^Cre^ 377 somata from 3 cultures). 1-way ANOVA with post hoc Tukey’s test. * p<0.05, ** p<0.01, ***p<0.001, ****p<0.0001.

**Supplemental Figure 17.**
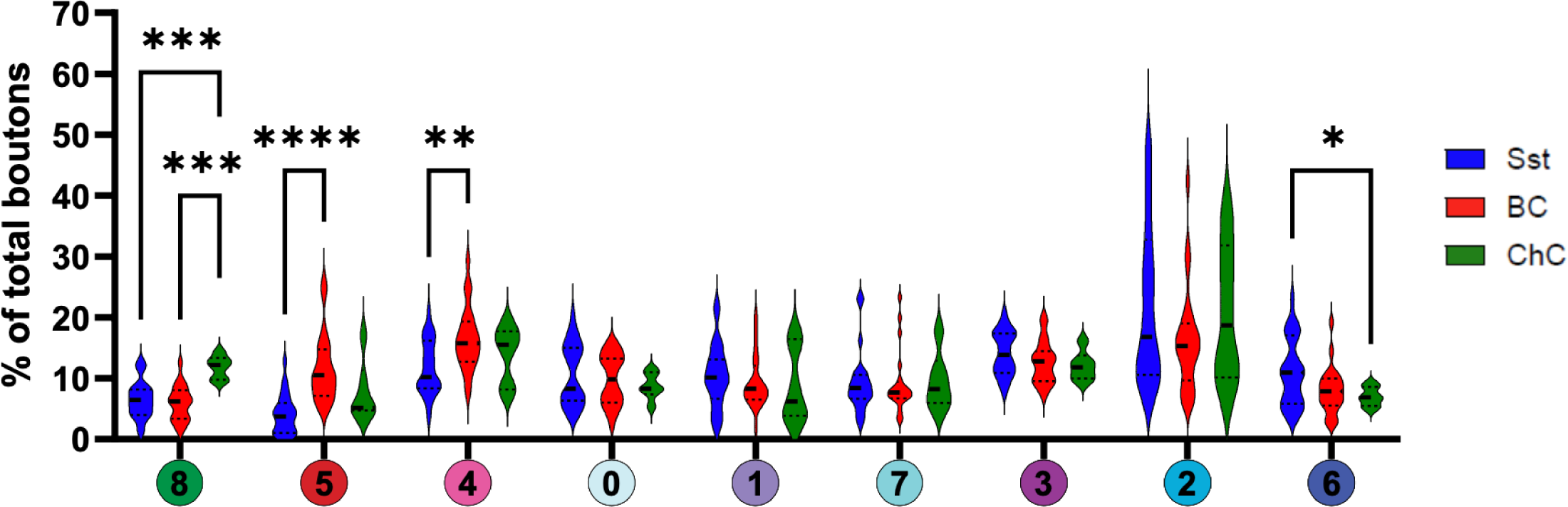
Relative proportion of synaptic subgroups employed by Sst, BCs and ChCs.

**Supplemental Figure 18.**
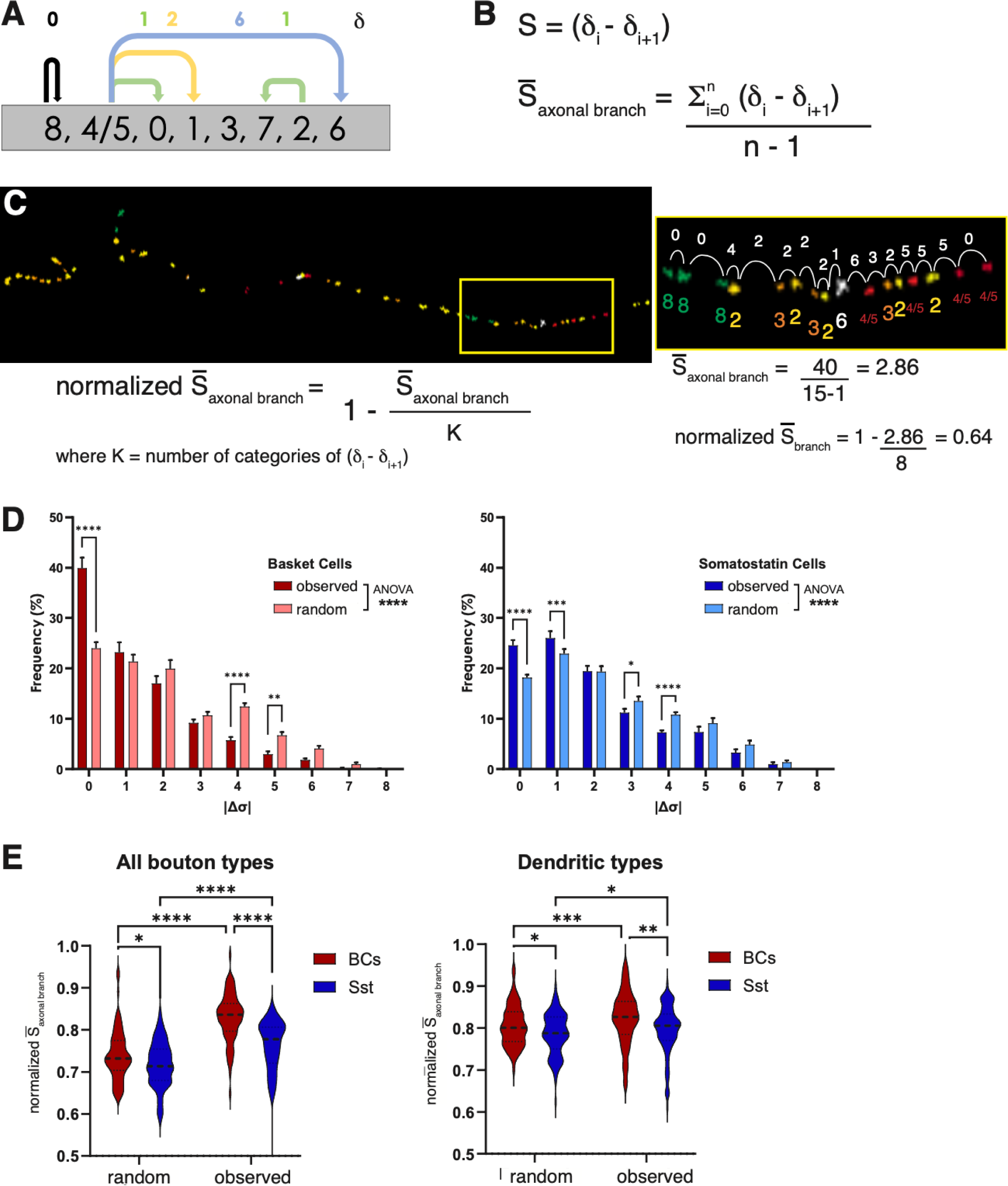
Calculation of sequentiality score of axonal branches. (A) Optimal subgroup sequence (as determined previously in Figure 4, Supplemental Figure 13) is used as the basis for calculating sequentiality score. The subgroup identity of synapses is compared to each successive synapse to count the ‘jump’ from optimal sequence. (B) Formula for calculating average sequentiality score for an axonal branch 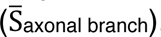. (C) Example of axonal branch with synaptic subgroup identity mapped onto original image. Box on the right is enlarge image of boxed region on left. White arcs depict comparison of subgroup identity across successive synapses and the ‘jump’ in sequentiality score. Bottom right shows calculation of sequentiality score for stretch of axonal branch evaluated. Next, 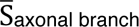 is normalized to a scale of 0 to 1 with 0 being the least sequential and 1 being the most. (D) Raw sequentiality data was randomized across 10,000 iterations to generate the random frequency distribution of average axonal branch sequentiality, 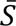. This was compared with observed 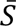 for both BCs (left) and Sst interneurons (right). (E) Randomized and observed 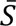 compared for all boutons (left) and for dendrite-targeting boutons alone. In both cases, observed boutons are significantly less random but more so for BCs than Sst interneurons.

**Supplemental Figure 19.** Summary of findings related to the spatial distribution of synaptic subgroups. (left) Schematic of interneuron subclasses forming canonical synapses onto a postsynaptic neuron. The panels on the right show how the model has changed with each successive finding: (1) We found dendrite targeting sub-groups that were spacially localized to proximal, mid, and distal subdomains of the dendrite, revealing a more complex postsynaptic topology. (2) Soma-targeting subgroups were distributed in different proportions across 3 groups of postsynaptic neurons that exhibited distinct features. (3) BCs and Sst interneurons enact inhibitory coverage via different strategies. BCs form numerous synapses along the postsynaptic neuron while Sst interneurons form fewer synapses before ‘skipping’ to another process.

